# Structure in the variability of the basic reproductive number (*R*_0_) for Zika epidemics in the Pacific islands

**DOI:** 10.1101/064949

**Authors:** Clara Champagne, David Georges Salthouse, Richard Paul, Van-Mai Cao-Lormeau, Benjamin Roche, Bernard Cazelles

## Abstract

Before the outbreak that reached the Americas in 2015, Zika virus (ZIKV) circulated in Asia and the Pacific: these past epidemics can be highly informative on the key parameters driving virus transmission, such as the basic reproduction number (*R*_0_). We compare two compartmental models with different mosquito representations, using surveillance and seroprevalence data for several ZIKV outbreaks in Pacific islands (Yap, Micronesia 2007, Tahiti and Moorea, French Polynesia 2013-2014, New Caledonia 2014). Models are estimated in a stochastic framework with state-of-the-art Bayesian techniques. *R*_0_ for the Pacific ZIKV epidemics is estimated between 1.5 and 4.1, the smallest islands displaying higher and more variable values. This relatively low range of *R*_0_ suggests that intervention strategies developed for other flaviviruses should enable as, if not more effective control of ZIKV. Our study also highlights the importance of seroprevalence data for precise quantitative analysis of pathogen propagation, to design prevention and control strategies.

---

In May 2015, the first local cases of Zika were recorded in Brazil and by December of the same year the number of cases had surpassed 1.5 million. On February 2016, the World Health Organization declared Zika as a public health emergency of international concern^1^ and in March 2016, local transmission of Zika was recognized in 34 countries. Previously the Zika virus had circulated in Africa and Asia but only sporadic human cases had been reported. In 2007 the outbreak on Yap (Micronesia) was the first Zika outbreak outside Africa and Asia.^2^ Since, Zika outbreaks have been also reported in French Polynesia and in New Caledonia^3;4^ between 2013 and 2014 and subsequently, there have been cases of Zika disease in the Cook Islands, the Solomon Islands, Samoa, Vanuatu, and Easter Island (Chile) (see Fig. 1 in reference^5^).

**Figure 1.**
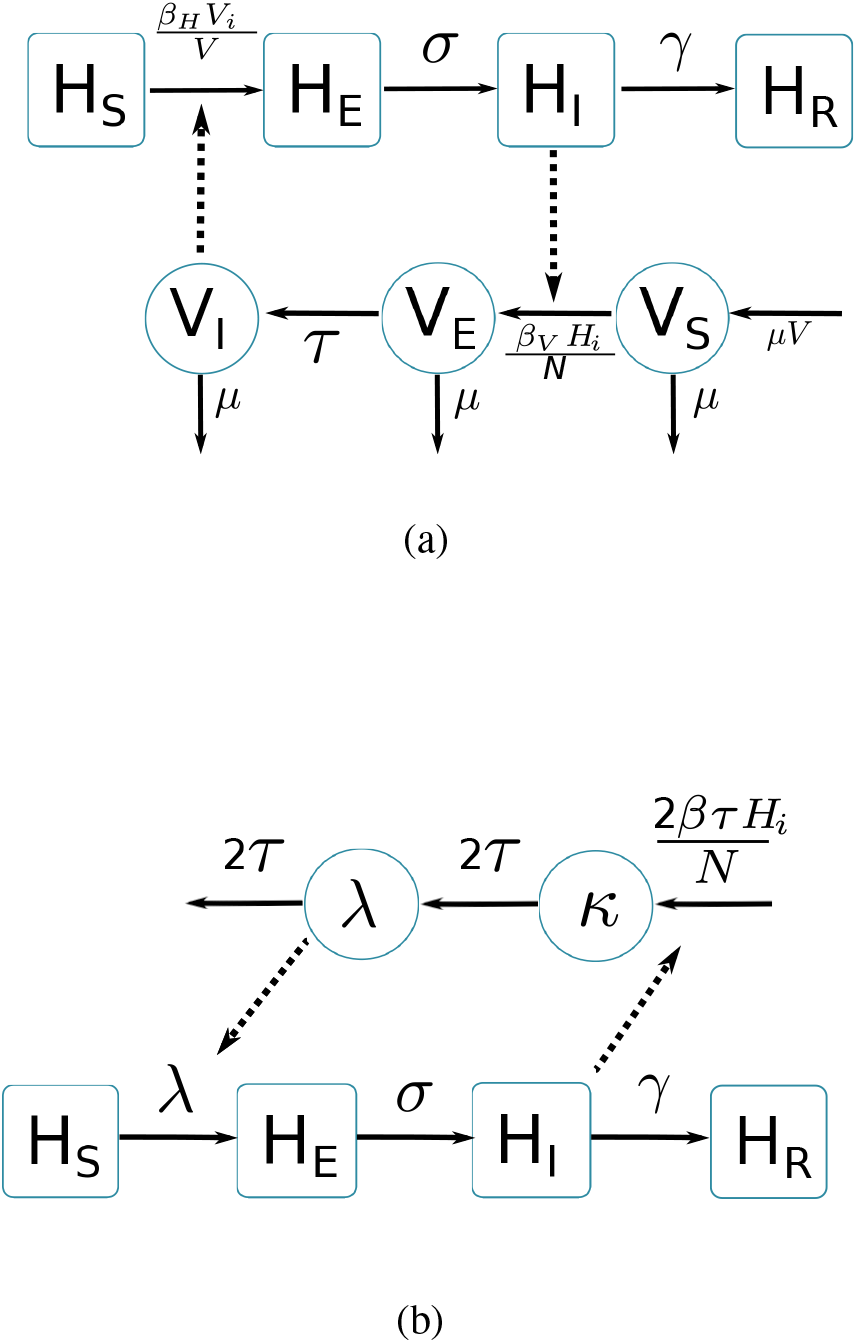
Graphical representation of compartmental models. Squared boxes and circles correspond respectively to human and vector compartments. Plain arrows represent transitions from one state to the next. Dashed arrows indicate interactions between humans and vectors. a) Pandey model.^16^ *H_S_* susceptible individuals; *H_E_* infected (not yet infectious) individuals; *H_I_* infectious individuals; *H_R_* recovered individuals; *σ* is the rate at which *H_E_*-individuals move to infectious class *H_I_*; infectious individuals (*H_I_*) then recover at rate *γ*; *V_S_* susceptible vectors; *V_E_* infected (not yet infectious) vectors; *V_I_* infectious vectors; *V* constant size of total mosquito population; *τ* is the rate at which *V_E_*-vectors move to infectious class *V_I_*; vectors die at rate *μ*. b) Laneri model.^17^ *H_S_* susceptible individuals; *H_E_* infected (not yet infectious) individuals; *H_I_* infectious individuals; *H_R_* recovered individuals; *σ* is the rate at which *H_E_*-individuals move to infectious class *H_I_*; infectious individuals (*H_I_*) then recover at rate *γ*; implicit vector-borne transmission is modeled with the compartments *κ* and *λ*; *λ* current force of infection; *κ* latent force of infection reflecting the exposed state for mosquitoes during the extrinsic incubation period; *τ* is the transition rate associated to the extrinsic incubation period.

Zika virus (ZIKV) is a flavivirus, mostly transmitted via the bites of infected *Aedes* mosquitoes, although non-vector-borne transmission has been documented (sexual and maternofoetal transmission, laboratory contamination, transmission through transfusion).^6^ The most common clinical manifestations include rash, fever, arthralgia, and conjunctivitis^6^ but a large proportion of infections are asymptomatic or trigger mild symptoms that can remain unnoticed. Nevertheless, the virus may be involved in many severe neurological complications, including Guillain-Barre syndrome^7^ and microcephaly in newborns.^8^ These complications and the impressive speed of its geographically propagation make the Zika pandemic a public health threat.^1^ This reinforces the urgent need to characterize the different facets of virus transmission and to evaluate its dispersal capacity. We address this here by estimating the key parameters of ZIKV transmission, including the basic reproduction number (*R*_0_), based on previous epidemics in the Pacific islands.

Defined as the average number of secondary cases caused by one typical infected individual in an entirely susceptible population, the basic reproduction number (*R*_0_) is a central parameter in epidemiology used to quantify the magnitude of ongoing outbreaks and it provides insight when designing control interventions.^9^ It is nevertheless complex to estimate^9;10^, and therefore, care must be taken when extrapolating the results obtained in a specific setting, using a specific mathematical model. In the present study, we explore the variability of *R*_0_ using two state-of-the-art models in several settings that had Zika epidemics in different years and that vary in population size (Yap, Micronesia 2007, Tahiti and Moorea, French Polynesia 2013-2014, and New Caledonia 2014). These three countries were successively affected by the virus, resulting in the first significant human outbreaks and they differ in several ways, including population size and location specific features. Hence, the comparison of their parameter estimates can be highly informative on the intrinsic variability of *R*_0_. For each setting, we compare two compartmental models using different parameters defining the mosquito populations. Both models are considered in a stochastic framework, a necessary layer of complexity given the small population size and state-of-the-art Bayesian inference techniques^11^ are used for parameter estimation.

## Results

We use mathematical transmission models and data from surveillance systems and seroprevalence surveys for several ZIKV outbreaks in Pacific islands (Yap, Micronesia 2007^2^, Tahiti and Moorea, French Polynesia 2013-2014^12–14^, New Caledonia 2014^15^) to quantify the ZIKV transmission variability.

Two compartmental models with vector-borne transmission are compared (cf. Figure 1). Both models use a Susceptible-Exposed-Infected-Resistant (SEIR) framework to describe the virus transmission in the human population, but differ in their representation of the mosquito population. Figure 1.a. is a schematic representation derived from Pandey et al.^16^ and formulates explicitly the mosquito population, with a Susceptible-Exposed-Infected (SEI) dynamic to account for the extrinsic incubation period (time taken for viral dissemination within the mosquito).

By contrast, in the second model (Figure 1.b.) based on Laneri et al.^17^ the vector is modeled implicitly: the two compartments *κ* and *λ* do not represent the mosquito population but the force of infection for vector to human transmission. This force of infection passes through two successive stages in order to include the delay associated with the extrinsic incubation period: *κ* stands for this latent phase of the force of infection whereas *λ* corresponds directly to the rate at which susceptible humans become infected.

The basic reproduction number of the models (*R*_0_) is calculated using the next Generation Matrix method:^9^

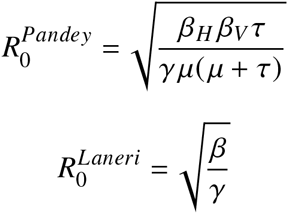

In addition, we consider that only a fraction *ρ* of the total population is involved in the epidemic, due to spatial heterogeneity, immuno-resistance, or cross-immunity. For both models we define *N* = *ρ* · *H* with H the total size of the population reported by census.

The dynamics of ZIKV transmission in these islands is highly influenced by several sources of uncertainties. In particular, the small population size (less than 7,000 inhabitants in Yap) leads to high variability in transmission rates. Therefore all these models are simulated in a discrete stochastic framework (Poisson with stochastic rates^18^), to take this phenomenon into account. Stochasticity requires specific inference techniques: thus estimations are performed with PMCMC algorithm (particle Markov Chain Monte Carlo^11^).

Using declared Zika cases from different settings, the two stochastic models (Fig. 1) were fitted (Figs 2-3). These models allow us to describe the course of the observed number of cases and estimate the number of secondary cases generated, *R*_0_. Our estimates of *R*_0_ lie between 1.6 (1.5-1.7) and 3.2 (2.4-4.1) and vary notably with respect to settings and models (Figures 2-3 and Tables 1-2). Strikingly, Yap displays consistently higher values of *R*_0_ in both models and in general, there is an inverse relationship between island size and both the value and variability of *R*_0_. This phenomenon may be explained by the higher stochasticity and extinction probability associated with smaller populations and can also reflect the information contained in the available data. However, the two highly connected islands in French Polynesia, Tahiti and Moorea, display similar values despite their differing sizes.

**Figure 2.**
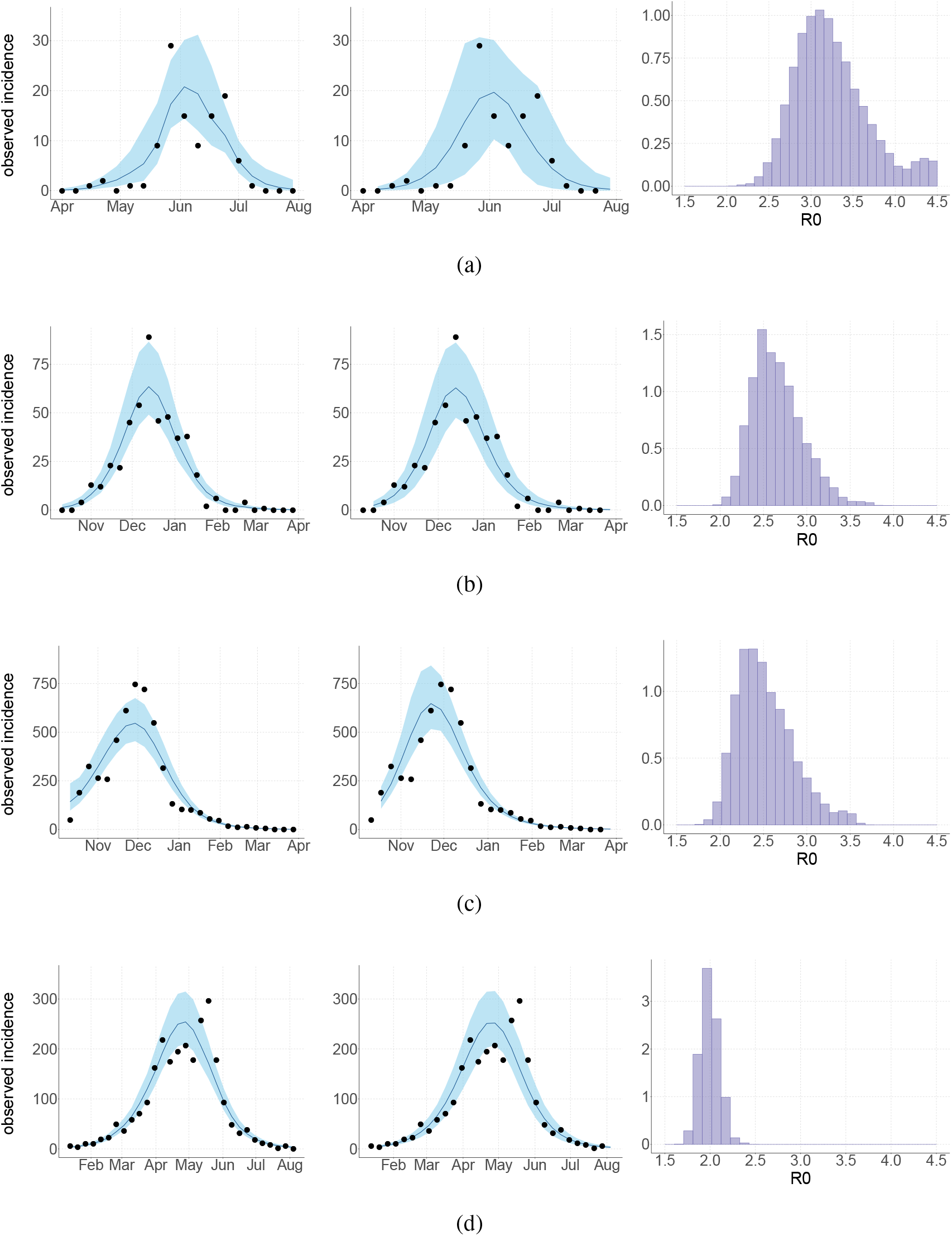
Results using the Pandey model. Posterior median number of observed Zika cases (solid line), 95% credible intervals (shaded blue area) and data points (black dots). First column: particle filter fit. Second column: Simulations from the posterior density. Third column: *R*_0_ posterior distribution. a) Yap. b) Moorea. c) Tahiti. d) New Caledonia. The estimated seroprevalences at the end of the epidemic (with 95% credibility intervals) are: a) 73% (CI95: 68-77, observed 73%); b) 49% (CI95: 45-53, observed 49%); c) 49% (CI95: 45-53, observed 49%); d) 39% (CI95: 8-92). See Figure 4.

**Figure 3.**
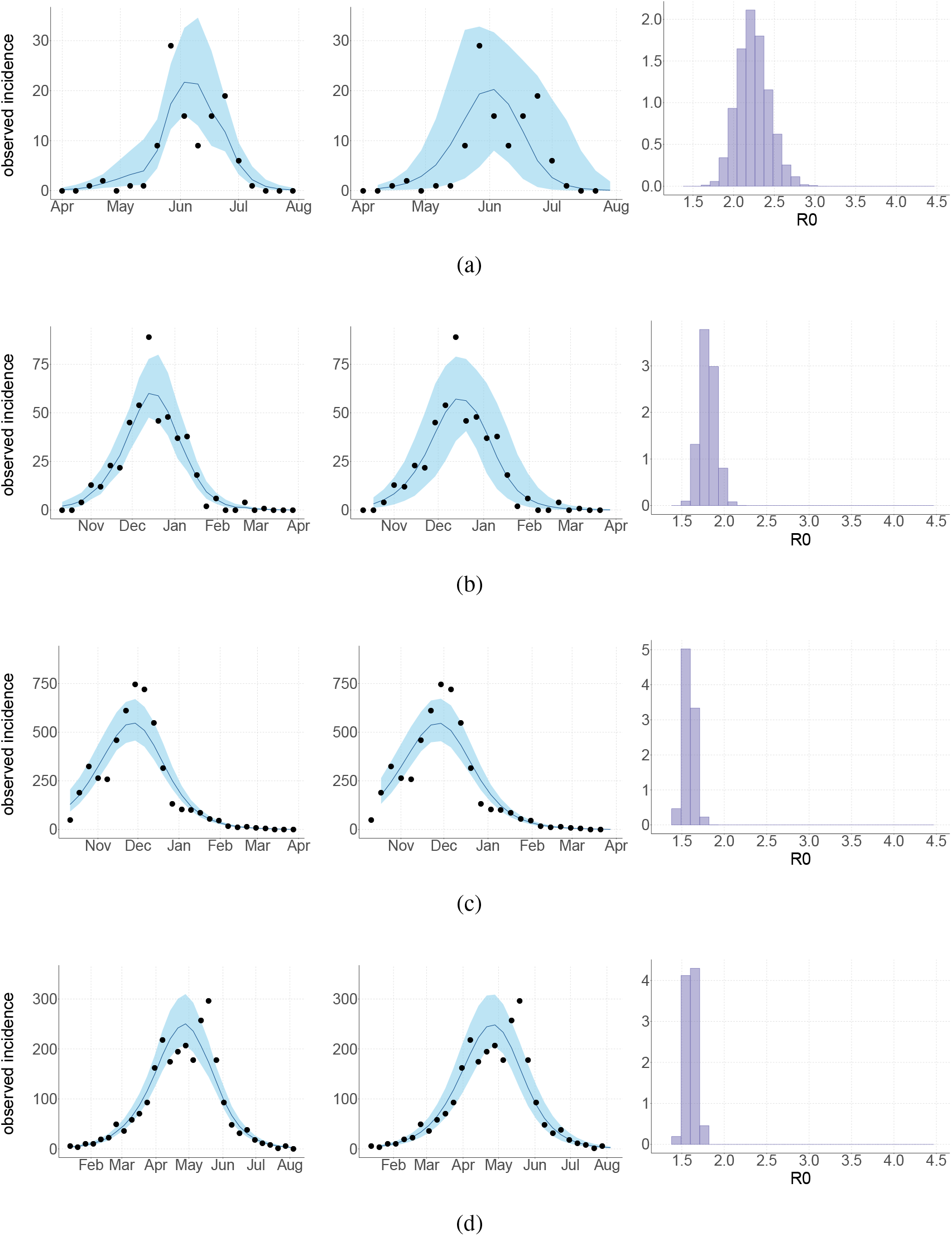
Results using the Laneri model. Posterior median number of observed Zika cases (solid line), 95% credible intervals (shaded blue area) and data points (black dots). First column: particle filter fit. Second column: Simulations from the posterior density. Third column: *R*_0_ posterior distribution. a) Yap. b) Moorea. c) Tahiti. d) New Caledonia. The estimated seroprevalences at the end of the epidemic (with 95% credibility intervals) are: a) 72% (CI95: 68-77, observed 73%); b) 49% (CI95: 45-53, observed 49%); c) 49% (CI95: 45-53, observed 49%); d) 65% (CI95: 24-91). See Figure 5.

Regarding model variability, *R*_0_ estimates are always higher and coarser with the Pandey model than with the Laneri model (cf. Tables 1-2). The Pandey model has two additional estimated parameters (in particular, the mosquito lifespan), which can explain the higher variability of the output. It is worth noting that these parameters are very sensitive (see Materials and methods). The difference in *R*_0_ may also be linked to the difference in the estimated initial number of infected individuals (*H*_*I*0_), which is higher in the Laneri model than in the Pandey model. Because of the high proportion of asymptomatic cases (the ratio of asymptomatic:symptomatic is estimated to be 1:1.3, V.-M Cao-Lormeau personal communication), it is hard to determine which scenario is more realistic, the time between introduction of the disease into the island and the first reported symptomatic case being unknown in most settings.

**Table 1.** Parameter estimations for the Pandey model. Posterior median (95% credible intervals). All the posterior parameter distributions are presented in Figures 6-9.

| PANDEY MODEL |  | Yap | Moorea | Tahiti | New Caledonia |
| --- | --- | --- | --- | --- | --- |
| Population size | $H$ | 6,892 | 16,200 | 178,100 | 268,767 |
| Basic reproduction number | $R_0$ | 3.2 (2.4 – 4.1) | 2.6 (2.2 – 3.3) | 2.5 (2.0 – 3.3) | 2.0 (1.8 – 2.2) |
| Observation rate | $r$ | 0.024 (0.018 – 0.031) | 0.058 (0.047 – 0.071) | 0.060 (0.048 – 0.072) | 0.029 (0.009 – 0.10) |
| Fraction of population involved | $\rho$ | 74% (69-79) | 50% (46-54) | 50% (46-54) | 34% (8-91) |
| Initial number of infected individuals | $H_{I0}$ | 2 (1-8) | 5 (0-21) | 304 (16-1145) | 37 (1-386) |
| Infectious period in human (days) | $\gamma^{-1}$ | 5.2 (4.0 – 6.5) | 5.2 (4.0 – 6.6) | 5.2 (4.0 – 6.5) | 5.4 (4.1 – 6.7) |
| Extrinsic incubation period in mosquito (days) | $\tau^{-1}$ | 10.5 (8.6 – 12.4) | 10.5 (8.6 – 12.4) | 10.4 (8.5 – 12.5) | 10.7 (8.9 – 12.5) |
| Mosquito lifespan (days) | $\mu^{-1}$ | 15.4 (11.8 – 19.3) | 15.3 (11.6 – 19.2) | 14.9 (10.9 – 18.8) | 15.3 (11.5 – 19.4) |

**Table 2.** Parameter estimations for the Laneri model. Posterior median (95% credible intervals). All the posterior parameter distributions are presented in Figures 10-13.

| LANERI MODEL |  | Yap | Moorea | Tahiti | New Caledonia |
| --- | --- | --- | --- | --- | --- |
| Population size | $H$ | 6,892 | 16,200 | 178,100 | 268,767 |
| Basic reproduction number | $R_0$ | 2.2 (1.9 – 2.6) | 1.8 (1.6 – 2.0) | 1.6 (1.5 – 1.7) | 1.6 (1.5 – 1.7) |
| Observation rate | $r$ | 0.02 (0.018 – 0.032) | 0.057 (0.047 – 0.071) | 0.057 (0.048 – 0.068) | 0.015 (0.009 – 0.033) |
| Fraction of population involved | $\rho$ | 73% (69 – 78) | 51% (46 – 54) | 54% (49 – 59) | 70% (31 – 100) |
| Initial number of infected individuals | $Hi_0$ | 2 (1 – 10) | 9 (1 – 28) | 667 (22 – 1570) | 82 (2 – 336) |
| Infectious period in human (days) | $\gamma^{-1}$ | 5.3 (4.0 – 6.6) | 5.2 (4.0 – 6.6) | 5.1 (4.0 – 6.5) | 5.4 (4.0 – 6.7) |
| Extrinsic incubation period in mosquito (days) | $\tau^{-1}$ | 10.6 (8.8 – 12.7) | 10.6 (8.6 – 12.6) | 10.5 (8.5 – 12.4) | 10.8 (8.9 – 12.7) |

For the duration of infectious and intrinsic incubation (in human) and extrinsic incubation (in mosquito) periods, the posterior density ressembles the informative prior (cf. Figures 6-13), indicating the models’ incapacity to identify properly these parameters without more informative data. Moreover, these parameters have a clear sensitivity (see Materials and methods) and precise field measures are therefore crucial for reliable model predictions.

The fraction *ρ* of the population involved in the epidemic and the observation rate *r* display very large credible intervals when seroprevalence is unknown (New Caledonia). They are highly correlated with one another (cf. Figures 17 and 21) and therefore unidentifiable without precise information on seroprevalence.

## Discussion

The reproduction number *R*_0_ is a key parameter in epidemiology that characterizes the epidemic dynamics and the initial spread of the pathogen at the start of an outbreak in a susceptible population. *R*_0_ can be used to inform public health authorities on the level of risk posed by an infectious disease, vaccination strategy, and the potential effects of control interventions^19^. In the light of the potential public health crisis generated by the international propagation of ZIKV, characterization of the potential transmissibility of this pathogen is crucial for predicting epidemic size, rate of spread and efficacy of intervention.

Using data from both surveillance systems and seroprevalence surveys in four different geographical settings across the Pacific,^2;12-15^ we have estimated the basic reproductive number *R*_0_ (see Figs 2-3 and Tables 1-2). Our estimate of *R*_0_ obtained by inference based on Particle MCMC^11^ has values in the range 1.6(1.5-1.7) - 3.2 (2.4-4.1). Our *R*_0_ estimates vary notably across settings. Lower and finer *R*_0_ values are found in larger islands. This phenomenon can at least in part be explained by large spatial heterogeneities and higher demographic stochasticity for islands with smaller populations, as well as the influence of stochasticity on biological and epidemiological processes linked to virus transmission. This phenomenon can also be specific to the selection of the studied island or can reflect a highly clustered geographical pattern, the global incidence curve being the smoothed overview of a collection of more explosive small size outbreaks. However, it is notable that the two French Polynesian islands yield similar estimates of *R*_0_ despite differing population sizes. Indeed, other important factors differ among French Polynesia, New Caledonia and Yap, such as the human genetic background and their immunological history linked to the circulation of others arboviruses. Moreover, whilst both New Caledonia and French Polynesia populations were infected by the same ZIKV lineage and transmitted by the same principle vector species, *Aedes aegypti*, the epidemic in Yap occurred much earlier with a different ZIKV lineage^20^ and vectored by a different mosquito species *Aedes hensilli*.^21^ In French Polynesia, the vector *Aedes polynesiensis* is also present and dominates in Moorea with higher densities than in Tahiti. Finally, different vector control measures have been conducted in the three countries.

To date, studies investigating Zika outbreaks in the Pacific have always estimated *R*_0_ using a deterministic framework. Using a similar version of the Pandey model in French Polynesia, Kucharski et al.^22^ estimated *R*_0_ between 1.6 and 2.3 (after scaling to square root for comparison) for Tahiti and between 1.8 and 2.9 in Moorea. These estimates are slightly lower and less variable than ours. This difference can be explained firstly by the chosen priors on mosquito parameters and secondly because our model includes demographic stochasticity. Moreover, they predicted a seroprevalence rate at the end of the epidemic of 95-97%, far from the 49% measured. In Yap island, a study^23^ used a very detailed deterministic mosquito model, and estimated an *R*_0_ for Zika between 2.9 and 8. In this case, our lower and less variable estimates may come from the fact that our model is more parsimonious in the number of uncertain parameters, especially concerning the mosquito population. Finally, a third study^24^ relied on another method for *R*_0_ calculation (based only on the early exponential growth rate of the epidemic) in French Polynesia as a whole and in Yap. Again, the obtained parameters are lower than ours in French Polynesia and higher in Yap. In all these studies a deterministic framework is used excluding the possibility of accounting for the high variability of biological and epidemiological processes exacerbated by the small size of the population. In these three studies, like in ours, it is worth noting that little insight is obtained regarding mosquito parameters. Posterior distribution mimics the chosen prior (cf. Figures 6-13). Both the simulation of the epidemics and the estimated *R*_0_ are highly sensitive to the choice of priors on mosquito parameters, for which precise field measures are rare.

In the absence of sufficient data, the modeling of mosquito-borne pathogen transmission is a difficult task due to non-linearity and non-stationarity of the involved processes.^25^ This work has then several limitations. First, our study is limited by the completeness and quality of the data, with regard to both incidence and seroprevalence, but, above all, by the scarcity of information available on mosquitoes. Incidence data is aggregated at the island scale and cannot disentangle the effects of geography and observation noise to explain bimodal curves observed in Yap and New Caledonia. Moreover, although all data came from national surveillance systems, we had very little information about the potential degree of under-reporting. Seroprevalence data were gathered from small sample sizes and were missing in New Caledonia, which leads to strong correlation between the observation rate and the fraction of the population involved in the epidemic. Because of the high proportion of asymptomatic or mildly symptomatic cases, the magnitude of the outbreaks is difficult to evaluate without precise seroprevalence data^26^ or detection of mild, asymptomatic or pre-symptomatic infections.^27^ Considering vectors, no demographic data were available and this partly explains the large variability of our *R*_0_ estimations.

Secondly, incidence and seroprevalence data were difficult to reconcile; the use of incidence data led to higher infection rates than those observed in the seroprevalence data. This difficulty has been overcome by considering that only a fraction of the population (*ρ*) is involved in the epidemic and then our model manages to reproduce the observed seroprevalence rate. This exposed fraction could be the result of spatial heterogeneity and high clustering of cases and transmission, as observed for dengue. Finer scale incidence and seroprevalence data would be useful to explore this. Another explanation for higher predicted than observed infection rates could be due to interaction with other flaviviruses. The Zika outbreak was concomitant with dengue outbreaks in French Polynesia^12;13^ and New Caledonia.^15^ Examples of coinfection have been reported^4^ but competition between these close pathogens may also have occurred. Finally, mathematical models with vectorial transmission may tend to estimate high attack rates, sometimes leading to a contradiction between observed incidence and observed seroprevalence. Assumptions on the proportionality between infected mosquitoes and the force of infection, as well as the density-dependence assumption in these models could be questioned. Indeed even if these assumptions are at the heart of the mathematical models of mosquito-borne pathogen transmission^28;29^ a recent review,^30^ and recent experimental results^31;32^ question these important points.

On a final note, the estimates of *R*_0_ for ZIKV are similar to but generally on the lower side of estimates made for two other flaviviruses of medical importance, dengue and Yellow Fever viruses^33-35^, even though caution is needed in the comparison of studies with differing models, methods and data sources. Interventions strategies developed for dengue should thus enable as, if not more effective control of ZIKV, with the caveat that ZIKV remains principally a mosquito-borne pathogen with little epidemiological significance of the sexual transmission route.

In conclusion, using state-of-the art stochastic modeling methods, we have been able to determine estimates of *R*_0_ for ZIKV with an unexpected relationship with population size. Further data from the current Zika epidemic in South America that is caused by the same lineage as French Polynesia will enable us to confirm this relationship. Our study highlights the importance of gathering seroprevalence data, especially for a virus that often leads to an asymptomatic outcome and it would provide a key component for precise quantitative analysis of pathogen propagation to enable improved planning and implemention of prevention and control strategies.

## Materials and methods

### Data

During the 2007 outbreak that struck Yap, 108 suspected or confirmed Zika cases (16 per 1,000 inhabitants) were reported by reviewing medical records and conducting prospective surveillance between April 1st and July 29th 2007.^2^ In French Polynesia, sentinel surveillance recorded more than 8,700 suspected cases (32 per 1,000 inhabitants) across the whole territory between October 2013 and April 2014.^12;13^ In New Caledonia, the first Zika case was imported from French Polynesia on 2013 November 12th. Approximately 2,500 cases (9 per 1,000 inhabitants) were reported through surveillance between January (first autochtonous case) and August 2014.^15^

For Yap and French Polynesia, the post-epidemic seroprevalence was assessed. In Yap, a household survey was conducted after the epidemic, yielding an infection rate in the island of 73%.^2^ In French Polynesia, three seroprevalence studies were conducted. The first one took place before the Zika outbreak, and concluded that most of the population was naive for Zika virus.^36^ The second seroprevalence survey was conducted between February and March 2014, at the end of the outbreak, and reported a seroprevalence rate around 49%.^14^ The third one concerned only schoolchildren in Tahiti and was therefore not included in the present study.

Demographic data on population size were based on censuses from Yap^2^, French Polynesia^37^, and New Caledonia.^38^

## Models and inference

### Model equations

Although the models are simulated in a stochastic framework, we present them with ordinary differential equations for clarity. The reactions involved in the stochastic models are the same as those governed by the deterministic equations, but the simulation process differs through the use of discrete compartments. It is described in the next section.

The equations describing Pandey model are:

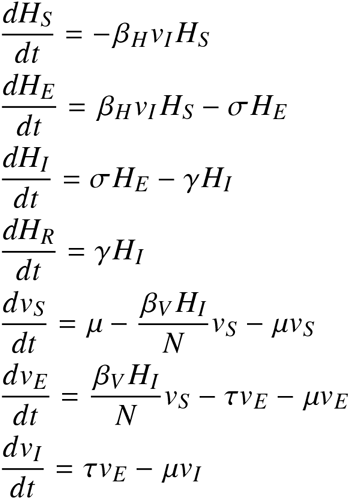

where 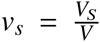 is the proportion of susceptible mosquitoes, 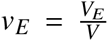 the proportion of exposed mosquitoes, and 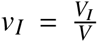 the proportion of infected mosquitoes. Since we are using a discrete model, we cannot use directly the proportions *v_S_, v_E_* and *v_I_* whose values are smaller than one. Therefore, we rescale using *V* = *H*, which leads to 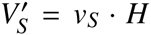, 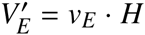, and 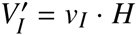.

In this model, the force of infection for humans is *λ_H_* = *β_H_v_I_*, and the force of infection for mosquitoes is 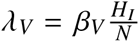

The equations describing Laneri model are:

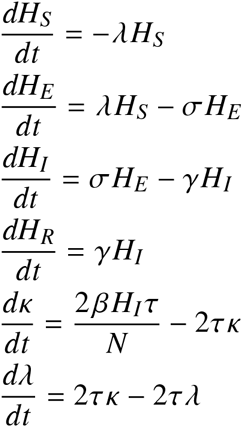

In this model, the role of mosquitoes in transmission is represented only through the delay they introduce during the extrinsic incubation period (EIP, incubation period in the mosquito). For modeling reasons, this delay is included by representing the force of infection from infected humans to susceptible humans with two compartments *κ* and *λ*: in this formalism, the duration between the moment when an exposed individual becomes infectious and the moment when another susceptible individual acquires the infection has a gamma distribution of mean *τ*^−1^.^17;39;40^ Therefore, *λ* represents the current force of infection for humans *λ_H_* = *λ*. The compartment *κ* represents the same force of infection but at a previous stage, reflecting the exposed phase for mosquitoes during the extrinsic incubation period. As an analogy to Pandey model, the force of infection for mosquitoes is 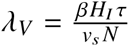, and therefore, the parameter *β* can be interpreted as the product of a transmission parameter *β*′ by the proportion of susceptible mosquitoes: *β* = *v_s_β*′. The force of infection for mosquitoes is then similar to Pandey’s: 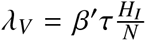.

Again, since we are using a discrete model, we cannot use directly the proportions *λ* and *κ* whose values are smaller than one. Therefore, we rescale up to a factor *N*, which leads to *L* = *λN* and *K* = *κN*.

### Stochastic framework

Both models are simulated in a stochastic and discrete framework, the Poisson with stochastic rates formulation,^18^ to include the uncertainties related to small population size. In this framework, the number of reactions occurring in a time interval *dt* is approximated by a multinomial distribution. In a model with *m* possible reactions and *c* compartments, *z_t_* being the state of the system at time *t* and *θ* the model parameters, the probability that each reaction *r^k^* occurs *n_k_* times in *dt* is calcutated as follows:^41^

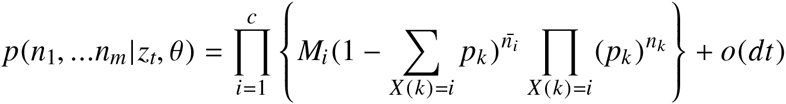

with, 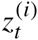 being the number of individual in compartment *i* at time *t*,

- 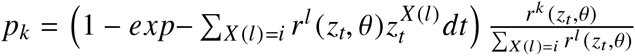
- 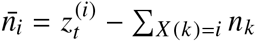 (the number of individuals staying in compartment *i* in *dt*)
- 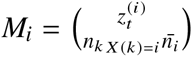 (multinomial coefficient)

### Observation models

The only observed compartments are the infected humans (incidence measured every week) and the recovered humans (seroprevalence at the end of the outbreak when data is available). In order to link the model to the data, two observation models, for both incidence and seroprevalence data, are needed.

### Observation model on incidence data

The observed weekly incidence is assumed to follow a negative binomial distribution^18^ whose mean equals the number of new cases predicted by the model times an estimated observation rate *r*.

The observation rate *r* accounts for non observed cases, due to non reporting from medical centers, mild symptoms unseen by health system, and asymptomatic infections. Without additional data, it is not possible to make a distinction between these three categories of cases. We also implicitely make the assumption that these cases transmit the disease as much as reported symptomatic cases.

The observation model for incidence data is therefore:

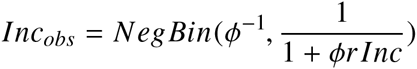

*Inc_obs_* being the observed incidence, and *Inc* the incidence predicted by the model. The dispersion parameter^18^ *ϕ* is fixed at 0.1.

### Observation model on seroprevalence data

Seroprevalence data is fitted for Tahiti, Moorea, and Yap settings. It is assumed that the observed seroprevalence at the end of the epidemic follows a normal distribution with fixed standard deviation, whose mean equals the number of individuals in the *H_R_* compartment predicted by the model.

The observation model for seroprevalence data is therefore:

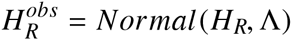

at the last time step, with notations detailed for each model in Table 3.

**Table 3.** Details of the observation models for seroprevalence.

| Island | Date | Standard deviation | Observed seroprevalence |
| --- | --- | --- | --- |
| | | $\Lambda$ | $H_R^{obs}$ |
| Yap | 2007-07-29 | 150 | 5005<sup>2</sup> |
| Moorea | 2014-03-28 | 325 | $0.49 \times 16200 = 7938$<sup>14</sup> |
| Tahiti | 2014-03-28 | 3562 | $0.49 \times 178100 = 87269$<sup>14</sup> |

### Prior distributions

Informative prior distributions are assumed for the mosquito lifespan, the duration of infectious period, and both intrinsic and extrinsic incubation periods. The initial numbers of infected mosquitoes and humans are estimated, and the initial number of exposed individuals is set to the initial number of infected to reduce parameter space. We assume that involved populations are naive to Zika virus prior to the epidemic and set the initial number of recovered humans to zero. The other priors and associated references are listed in Table 4.

**Table 4.** Prior distributions of parameters. “Uniform[0,20]” indicates a uniform distribution between in the range [0,20]. “Normal(5,1) in [4,7]” indicates a normal distribution with mean 5 and standard deviation 1, restricted to the range [4,7].

| Parameters |  | Pandey model |  | Laneri model |  | References |
| --- | --- | --- | --- | --- | --- | --- |
| $R_0^2$ | squared basic reproduction number | Uniform[0, 20] | | Uniform[0, 20] | | assumed |
| $\beta_V$ | transmission from human to mosquito | Uniform[0, 10] | | . | | assumed |
| $\gamma^{-1}$ | infectious period (days) | Normal(5, 1) in [4, 7] | | Normal(5, 1) in [4, 7] | | 13 |
| $\sigma^{-1}$ | intrinsic incubation period (days) | Normal(4, 1) in [2, 7] | | Normal(4, 1) in [2, 7] | | 42–44 |
| $\tau^{-1}$ | extrinsic incubation period (days) | Normal(10.5, 1) in [4, 20] | | Normal(10.5, 1) in [4, 20] | | 45;46 |
| $\mu^{-1}$ | mosquito lifespan (days) | Normal(15, 2) in [4, 30] | | . | | 47;48 |
| $\rho$ | fraction of population involved | Uniform[0, 1] | | Uniform[0, 1] | | |
| Initial conditions (t=0) |  | Pandey model |  | Laneri model |  |  |
| $H_{I\ 0}$ | Infected humans | Uniform[$10^{-6}$, 1]·N | | Uniform[$10^{-6}$, 1]·N | | |
| $H_{E\ 0}$ | exposed humans | $H_{I\ 0}$ | | $H_{I\ 0}$ | | |
| $H_{R\ 0}$ | Recovered humans | 0 | | 0 | | |
| | Infected vectors | $V_{I0}$=Uniform[$10^{-6}$, 1]·H | | $L_0$=Uniform[$10^{-6}$, 1]·N | | |
| | exposed vectors | $V_{E\ 0} = V_{I0}$ | | $K_0=L_0$ | | |
| Local conditions |  | Yap | Moorea | Tahiti | New Caledonia | References |
| r | Observation rate | Uniform[0, 1] | Uniform[0, 1] | Uniform[0, 0.3] | Uniform[0, 0.23] | 13;15;49 |
| H | Population size | 6,892 | 16,200 | 178,100 | 268,767 | 2;37;38 |

The range for the prior on observation rate is reduced for Tahiti and New Caledonia, in order to reduce the parameter space and facilitate convergence. In both cases, we use the information provided with the data source. In French Polynesia, 8,750 cases we reported, but according to local health authorities, more than 32,000 people would have attended health facilities for Zika^13^ (8750/32000 ≤ 0.3). In New Caledonia, approximately 2,500 cases were reported but more than 11,000 cases were estimated by heath authorities^49^ (2500/11000 ≤ 0.23). In both cases, these extrapolations are lower bounds on the real number of cases (in particular, they do not estimate the number of asymptomatic infections), and therefore can be used as upper bounds on the observation rate.

## Estimations

### Inference with PMCMC

The complete model is represented using the state space framework, with two equation systems: the transition equations refer to the transmission models, and the measurement equations are given by the observation models.

In a deterministic framework, this model could be directly estimated using MCMC, with a Metropolis-Hastings algorithm targeting the posterior distribution of the parameters. This algorithm would require the calculation of the model likelihood at each iteration.

In our stochastic framework, the model output is given only through simulations and the likelihood is intractable. In consequence, estimations are performed with the PMCMC algorithm (particle Markov Chain Monte Carlo^11^), in the PMMH version (particle marginal Metropolis-Hastings). This algorithm uses the Metropolis-Hastings structure, but replaces the real likelihood by its estimation with Sequential Monte Carlo (SMC).

#### Algorithm 1 PMCMC^11^ (PMMH version, as in SSM^41^)

~~~
In a model with *n* observations and *J* particles.
*q*(.|*θ*^(*i*)^) is the transition kernel.
1: Initialize *θ*^(0)^.
2: Using SMC algorithm, compute 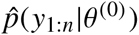 and sample 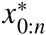 from 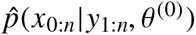.
3: **for** *i* = 1…*N^θ^* **do**
4:    Sample *θ*\* from *q*(.|*θ*^(*i*)^)
5:    Using SMC algorithm, compute 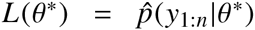 and sample 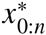 from 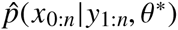
6:    Accept *θ*\* (et 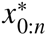) with probability 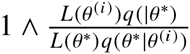
7:    If accepted, *θ*^(*i*+1)^ = *θ*\* and 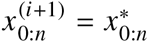. Otherwise *θ*^(*i*+1)^ = *θ*^(*i*)^ and 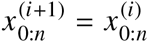.
8: **end for**
~~~

SMC^50^ is a filtering method that enables to recover the latent variables and estimate the likelihood for a given set of parameters. The data is treated sequentially, by adding one more data point at each iteration. The initial distribution of the state variables is approximated by a sample a particles, and from one iteration to the next, all the particles are projected according to the dynamic given by the model. The particles receive a weight according to the quality of their prediction regarding the observations. Before the next iteration, all the particles are resampled using these weights, in order to eliminate low weight particles and concentrate the computational effort in high probability regions. Model likelihood is also computed sequentially at each iteration^41;51^.

#### Algorithm 2 SMC (*Sequential Monte Carlo*, as implemented in SSM^41^)

~~~
In a model with *n* observations and *J* particles.
*L* is the model likelihood *p*(*y*_1:*T*_|*θ*). 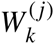 is the weight and 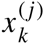 is the state associated to particle *j* at iteration *k*.
1: Set *L* = 1, 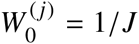.
2: Sample 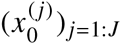 from *p*(*x*|*θ*_0_).
3: **for** *k* = 0: *n* − 1 **do**
4:   **for** *j* = 0: *J* **do**
5:          Sample 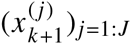 from *p*(*x*_*k*+1_|*x_k_, θ*)
6:          Set 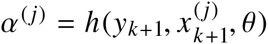
7:   **end for**
8:   Set 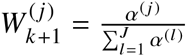 and 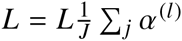
9:   Resample 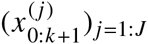 from 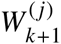
10: **end for**
~~~

A gaussian kernel *q*(.|*θ*^(*i*)^) is used in the PMCMC algorithm, with mean *θ*^(*i*)^ and fixed variance Σ^*q*^ (random walk Metropolis Hastings).

### Initialization

PMCMC algorithm is very sensitive to initialization of both the parameter values *θ*^(0)^ and the covariance matrix Σ^*q*^. Several steps of initialization are therefore used.

Firstly, parameter values are initialized by maximum likelihood through simplex algorithm on a deterministic version of the model. We apply the simplex algorithm to a set of 1000 points sampled in the prior distributions and we select the parameter set with the highest likelihood.

Secondly, in order to initialise the covariance matrix, an adaptative MCMC (Metropolis Hastings) framework is used^41;52^. It uses the empirical covariance of the chain Σ^(*i*)^, and aims to calibrate the acceptance rate of the algorithm to an optimal value. The transition kernel is also mixed (with a probability *α* = 0.05) with another gaussian using the identity matrix to improve mixing properties.

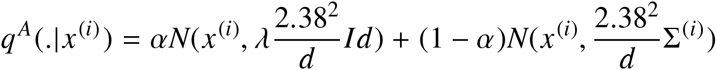

The parameter *λ* is approximated by successive iterations using the empirical acceptance rate of the chain.

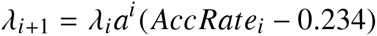

The adaptative PMCMC algorithm itself may have poor mixing properties without initialization. A first estimation of the covariance matrix is computed using KMCMC algorithm.^41^ In the KMCMC algorithm, the model is simulated with stochastic differential equations (intermediate between deterministic and Poisson with stochastic rates frameworks) and the SMC part of the adaptative PMCMC is replaced by the extended Kalman filter. When convergence is reached with KMCMC, then, adaptative PMCMC is used.

The PMCMC algorithm is finally applied on the output of the adaptative PMCMC, using 50,000 iterations and 10,000 particles. Calculations are performed with SSM software^41^ and R version 3.2.2.

## *R*_0_ calculation

*R*_0_ is calculated using the Next Generation Matrix approach^9^ (NGM).

### *R*_0_ calculation in Pandey model

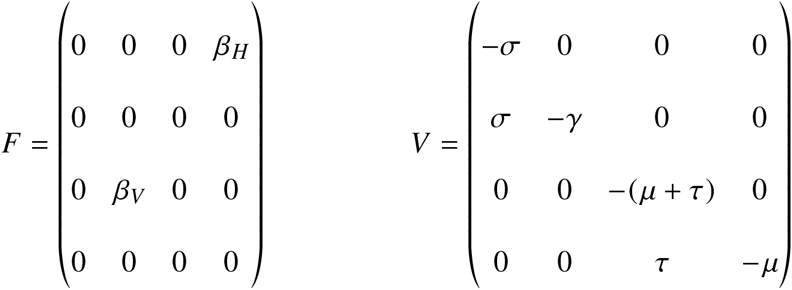

Then we have,

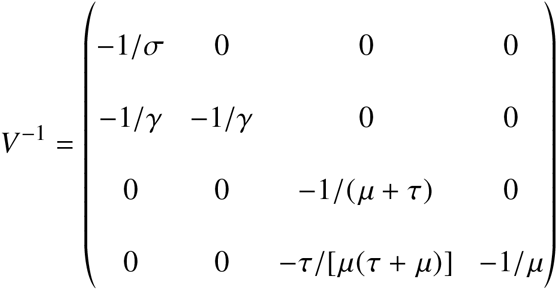

and

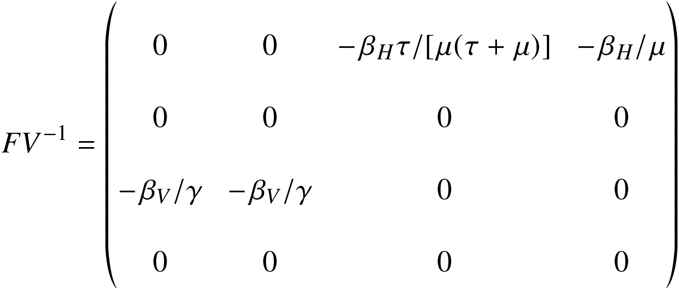

We calculate the eigen values *α* of −*FV*^−1^:

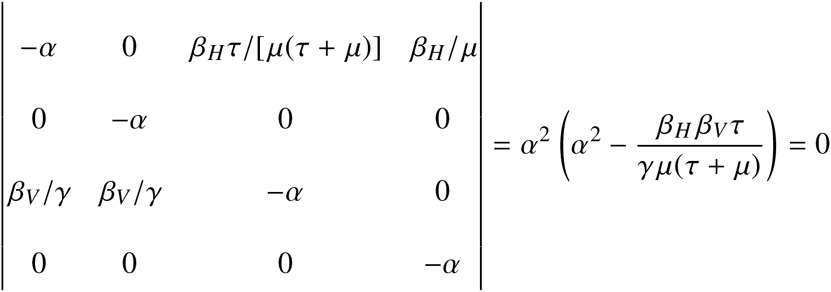

Then *α* = 0 or 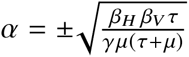 and the highest eigen value is 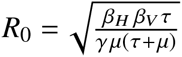.

This formula defines *R*_0_ as “the number of secondary cases per generation”^53^: i.e *R*_0_ can be written as the geometric mean 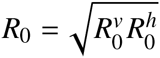, where 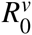 is the number of infected mosquitoes after the introduction of one infected human in a naive population, and 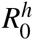 is the number of infected humans after the introduction of one infected mosquito in a naive population. With this definition, herd immunity is reached when 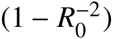 of the population is vaccinated^53^.

### *R*_0_ calculation in Laneri model

Following the analogy with Pandey model, we compute the spectral radius of the NGM for the Laneri model.

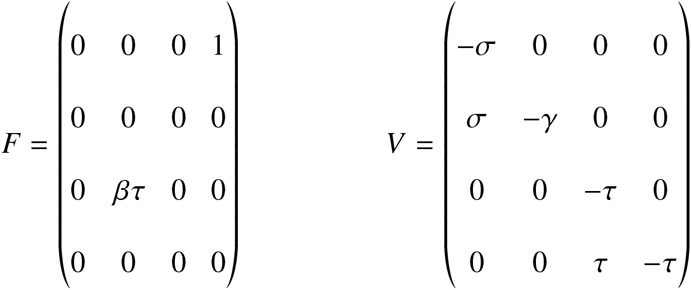

Then we have,

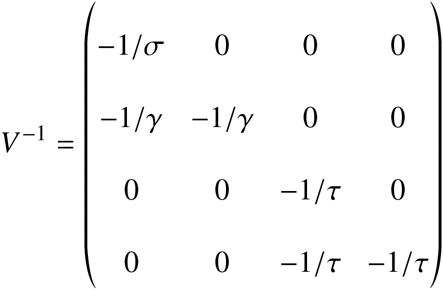

and

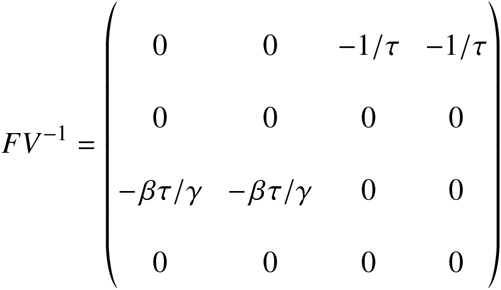

We calculate the eigen values *α* of −*FV*^−1^:

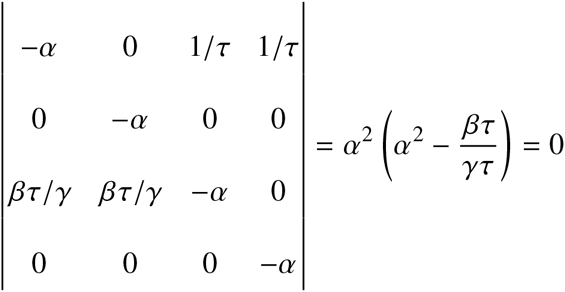

Then *α* = 0 or 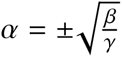 and the highest eigen value is 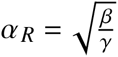.

Because *λ* and *κ* can be seen as parameters rather than state variables, the interpretation of the spectral radius as the *R*_0_ of the model is not straightforward. Therefore, we computed the *R*_0_ of the model through simulations, by counting the number of secondary infections after the introduction of a single infected individual in a naive population. Since Laneri model is considered here as a vector model, the number of infected humans after the introduction of a single infected is considered as 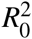. We simulated 1000 deterministic trajectories, using parameter values sampled in the posterior distributions for all parameters except initial conditions. With this method, the confidence intervals for number of infected humans 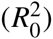 are similar to the ones of 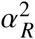 estimated by the model. As a consequence, *R*_0_ was approximated by the spectral radius of the NGM in our results with our stochastic framework (cf. Table 5).

**Table 5.** Square root of the number of secondary cases after the introduction of a single infected individual in a naive population. Median and 95% credible intervals of 1000 deterministic simulations using parameters from the posterior distribution.

|  | Pandey model | Laneri model |
| --- | --- | --- |
| Yap | 3.1 (2.5-4.3) | 2.2 (1.9-2.6) |
| Moorea | 2.6 (2.2-3.3) | 1.8 (1.6-2.0) |
| Tahiti | 2.5 (2.0-3.3) | 1.6 (1.5-1.7) |
| New Caledonia | 2.0 (1.8-2.2) | 1.6 (1.5-1.7) |

As a robustness check, the same method was applied to Pandey model: the confidence intervals for the number of secondary cases in simulations are very similar to the ones of 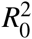 (cf. Table 5).

## Sensitivity analysis

In order to analyse the influence of parameter values on the model’s outputs, a sensitivity analysis was performed, using LHS/PRCC technique^54^, on Tahiti example. Similar results were obtained for the other settings. Three criteria were retained as outputs for the analysis: the seroprevalence at the last time point, the intensity of the peak of the outbreak and the date of the peak. We used uniform distributions for all parameters, which are listed in Tables 6 and 7. For model parameters, we used the same range as for the prior distribution. For initial conditions, the observation rate r and the fraction involved in the epidemic *ρ*, we used the 95% confidence interval obtained by PMCMC, in order to avoid unrealistic scenarios.

**Table 6.** Sensitivity analysis in Pandey model. Tahiti island. 1000 parameter sets were sampled with latin hypercube sampling (LHS), using “lhs” R package^55^. On each parameter set, the model was simulated deterministically. PRCC were computed using the “sensitivity” R package.^56^ P-values were calculated using the Student distribution approximation provided by Blower and Dowlatabadi.^54^

| Parameters |  | Seroprevalence |  | Peak intensity |  | Peak date |  |
| --- | --- | --- | --- | --- | --- | --- | --- |
|  | Distribution | PRCC | p-value | PRCC | p-value | PRCC | p-value |
| Model parameters |  |  |  |  |  |  |  |
| $R_0^2$ | Uniform[0,20] | 0.88 | <0.001 | 0.91 | <0.001 | -0.53 | <0.001 |
| $\beta_V$ | Uniform[0.1,10] | -0.69 | <0.001 | -0.77 | <0.001 | 0.29 | <0.001 |
| $\gamma^{-1}$ | Uniform[4,7] | -0.25 | <0.001 | 0.09 | 0.003 | 0.20 | <0.001 |
| $\sigma^{-1}$ | Uniform[2,7] | -0.03 | 0.324 | -0.12 | <0.001 | 0.18 | <0.001 |
| $\tau^{-1}$ | Uniform[4,20] | 0.00 | 0.921 | -0.05 | 0.102 | 0.08 | 0.012 |
| $\mu^{-1}$ | Uniform[4,30] | -0.61 | <0.001 | -0.75 | <0.001 | 0.51 | <0.001 |
| Initial conditions |  |  |  |  |  |  |  |
| $H_{i0}$ | Uniform[$2 \cdot 10^{-5}$,0.011] | -0.03 | 0.403 | -0.04 | 0.204 | 0.03 | 0.417 |
| $V_{i0}$ | Uniform[$10^{-4}$,0.034] | 0.06 | 0.044 | -0.03 | 0.290 | -0.24 | <0.001 |
| Fraction involved and observation model |  |  |  |  |  |  |  |
| $\rho$ | Uniform[0.46,0.54] | 0.50 | <0.001 | 0.14 | <0.001 | -0.03 | 0.307 |
| $r$ | Uniform[0.048,0.072] | 0.02 | 0.578 | 0.03 | 0.383 | 0.04 | 0.250 |

**Table 7.** Sensitivity analysis in Laneri model. Tahiti island. 1000 parameter sets were sampled with latin hypercube sampling (LHS), using “lhs” R package^55^. On each parameter set, the model was simulated deterministically. PRCC were computed using the “sensitivity” R package.^56^ P-values were calculated using the Student distribution approximation provided by Blower and Dowlatabadi.^54^

| Parameters |  | Seroprevalence |  | Peak intensity |  | Peak date |  |
| --- | --- | --- | --- | --- | --- | --- | --- |
|  | Distribution | PRCC | p-value | PRCC | p-value | PRCC | p-value |
| Model parameters |  |  |  |  |  |  |  |
| $R_0^2$ | Uniform[0,20] | 0.62 | <0.001 | 0.93 | <0.001 | -0.50 | <0.001 |
| $\gamma^{-1}$ | Uniform[4,7] | 0.01 | 0.731 | 0.62 | <0.001 | 0.15 | <0.001 |
| $\sigma^{-1}$ | Uniform[2,7] | -0.03 | 0.373 | -0.54 | <0.001 | 0.21 | <0.001 |
| $\tau^{-1}$ | Uniform[4,20] | -0.03 | 0.323 | -0.70 | <0.001 | 0.47 | <0.001 |
| Initial conditions |  |  |  |  |  |  |  |
| Hi<sub>0</sub> | Uniform[10<sup>-5</sup>,0.015] | 0.05 | 0.135 | 0.02 | 0.622 | -0.32 | <0.001 |
| L<sub>0</sub> | Uniform[2.10<sup>-5</sup>,0.004] | 0.05 | 0.133 | 0.00 | 0.930 | -0.16 | <0.001 |
| Fraction involved and observation model |  |  |  |  |  |  |  |
| $\rho$ | Uniform[0.49,0.59] | 0.80 | <0.001 | 0.34 | <0.001 | 0.02 | 0.558 |
| $r$ | Uniform[0.048,0.068] | -0.01 | 0.727 | 0.01 | 0.738 | -0.02 | 0.635 |

For all criteria, the key parameters in both models are transmission parameters (*R*_0_ and *β_V_*). High values for *R*_0_ are positively correlated with a large seroprevalence and a high and early peak. On the contrary, high values for the parameters introducing a delay in the model, the incubation periods in human (*σ*^−1^) and in mosquito (*τ*^−1^), are associated with a lower and later peak, and have no significant effect on seroprevalence. Moreover, the simulations are clearly sensitive to the other model parameters, in particular the mosquito lifespan (*μ*^−1^) in Pandey model.

Concerning other parameters, the initial conditions have a noticeable effect on the date of the peak only. As expected, the fraction involved in the epidemic (*ρ*) influences the magnitude of the outbreak, by calibrating the proportion of people than can be infected, but it has no significant effect on the timing of the peak.

## Complementary results

These complementary results include PMCMC results for both models in the four settings: the epidemic trajectories regarding the human compartments for infected and recovered individuals (Figures 4-5), the detailed posterior distributions for all parameters (Figures 6-13) and correlation plots for all models (Figures 14-21).

### Infected and recovered

**Figure 4.**
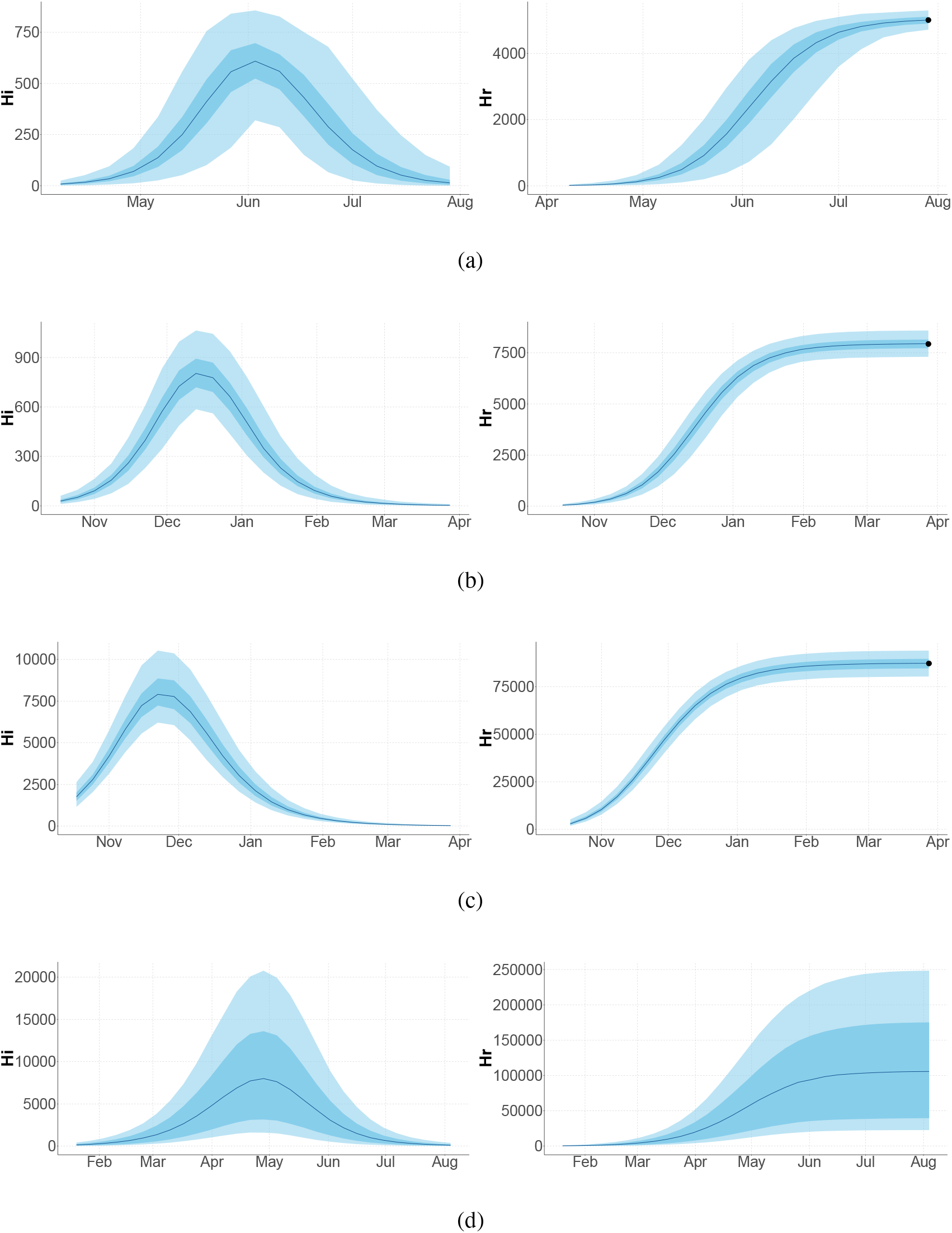
Infected and recovered humans evolution during the outbreak with Pandey model. Simulations from the posterior density: posterior median (solid line), 95% and 50% credible intervals (shaded blue areas) and observed seroprevalence (black dots). First column: Infected humans (*H_I_*). Second column: Recovered humans (*H_R_*). a) Yap. b) Moorea. c) Tahiti. d) New Caledonia.

**Figure 5.**
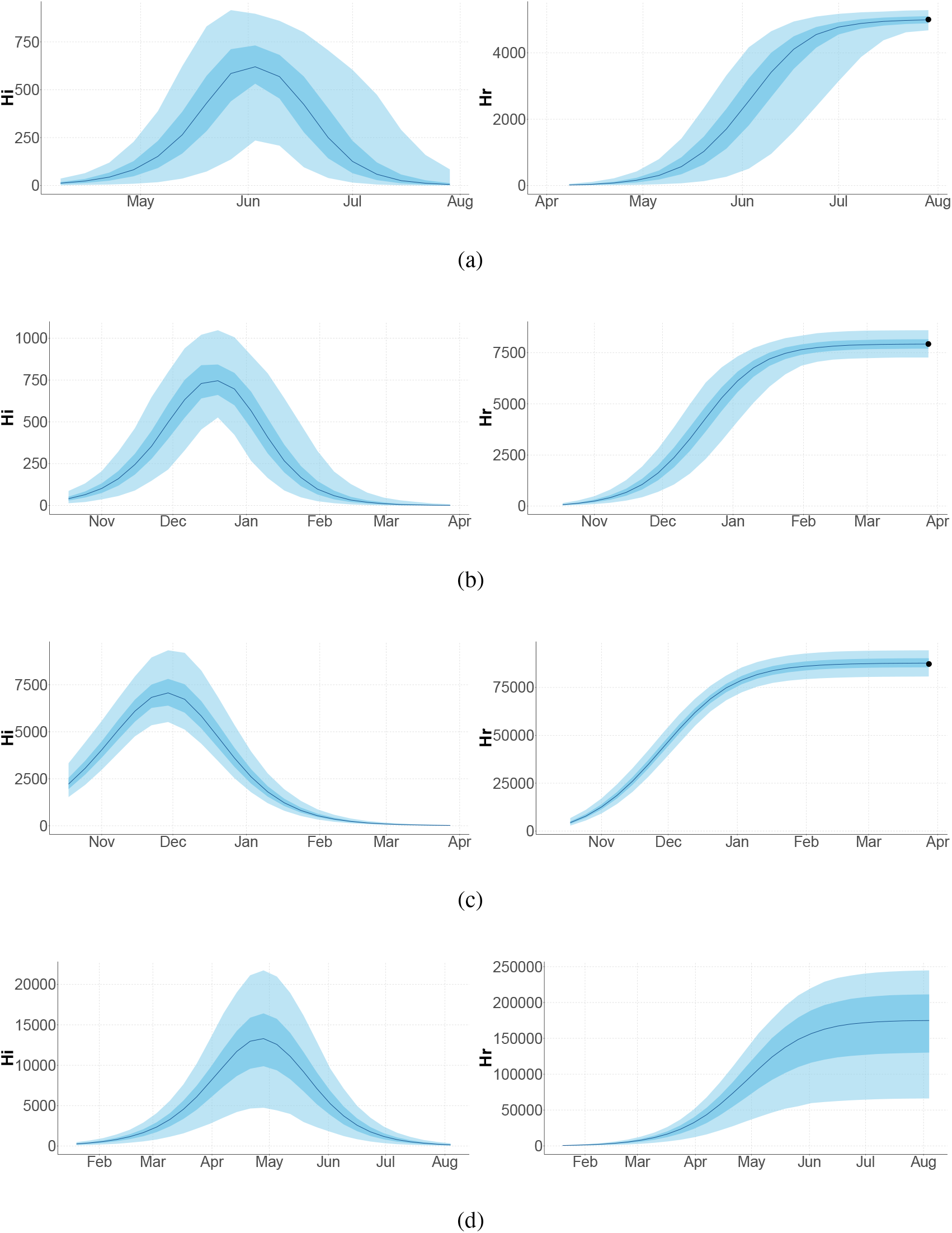
Infected and recovered humans evolution during the outbreak with Laneri model. Simulations from the posterior density: posterior median (solid line), 95% and 50% credible intervals (shaded blue areas) and observed seroprevalence (black dots). First column: Infected humans (*H_I_*). Second column: Recovered humans (*H_R_*). a) Yap. b) Moorea. c) Tahiti. d) New Caledonia.

### Posterior distributions

**Figure 6.**
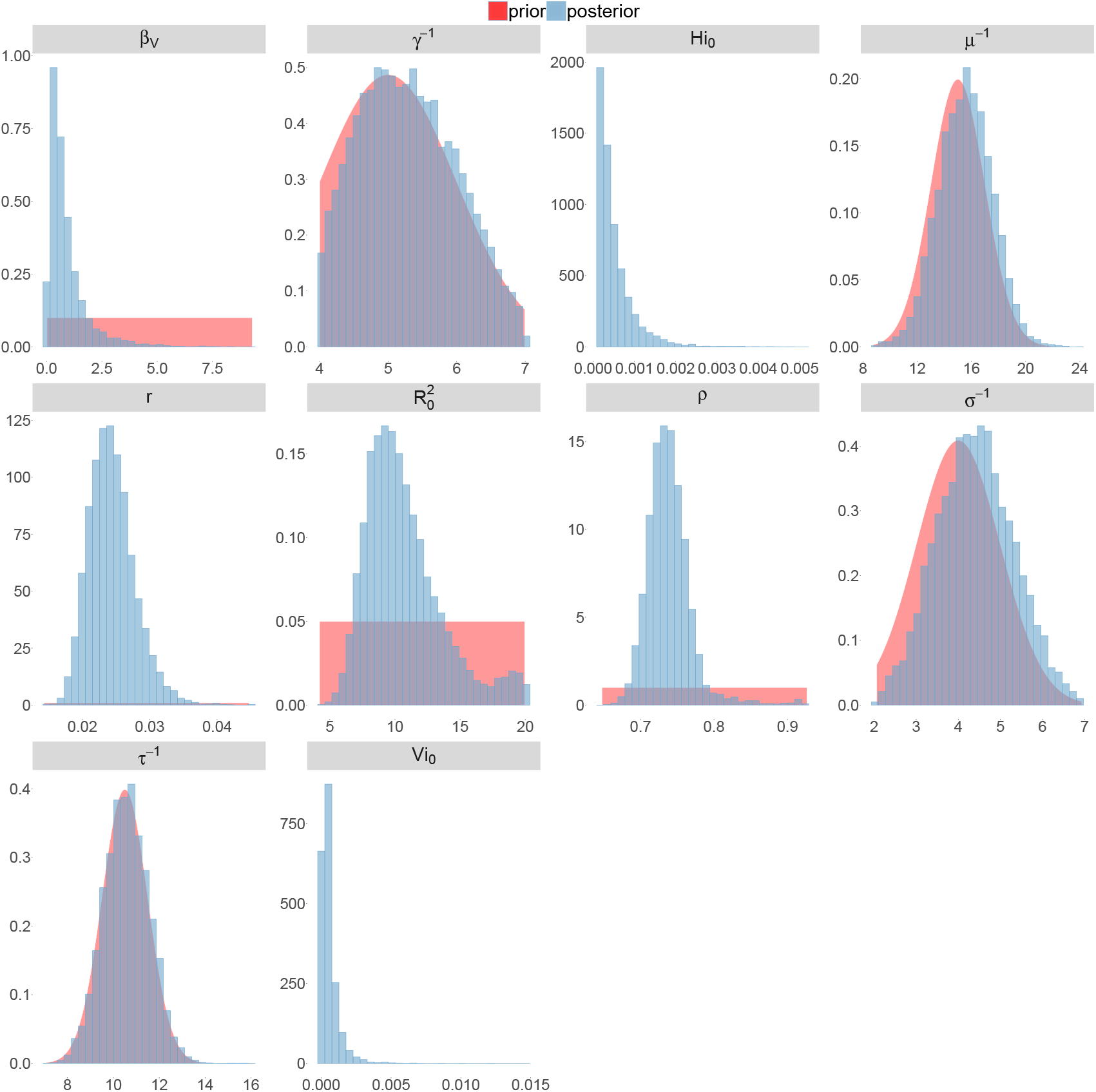
Posterior distributions. Pandey model, Yap island.

**Figure 7.**
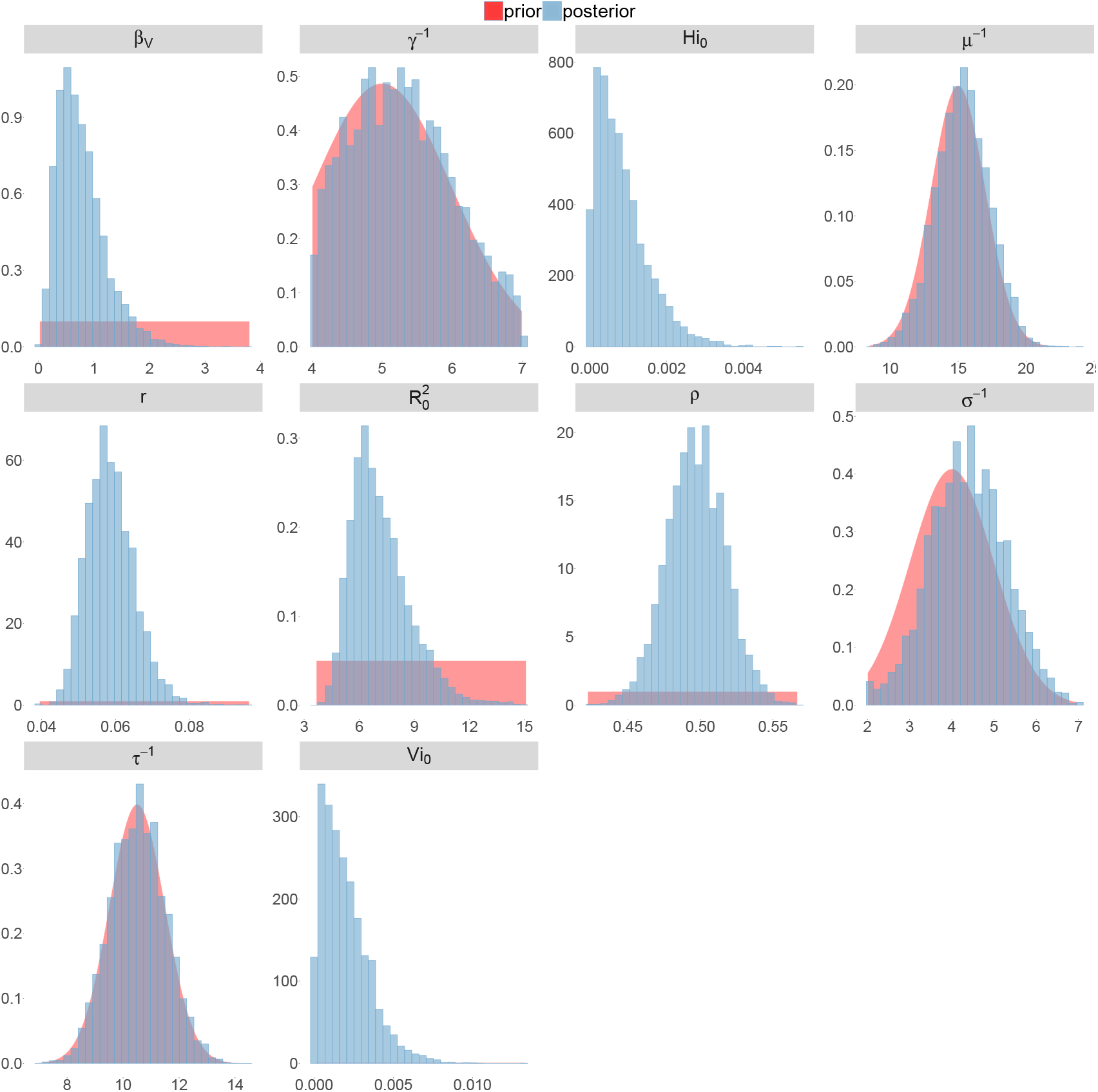
Posterior distributions. Pandey model, Moorea island.

**Figure 8.**
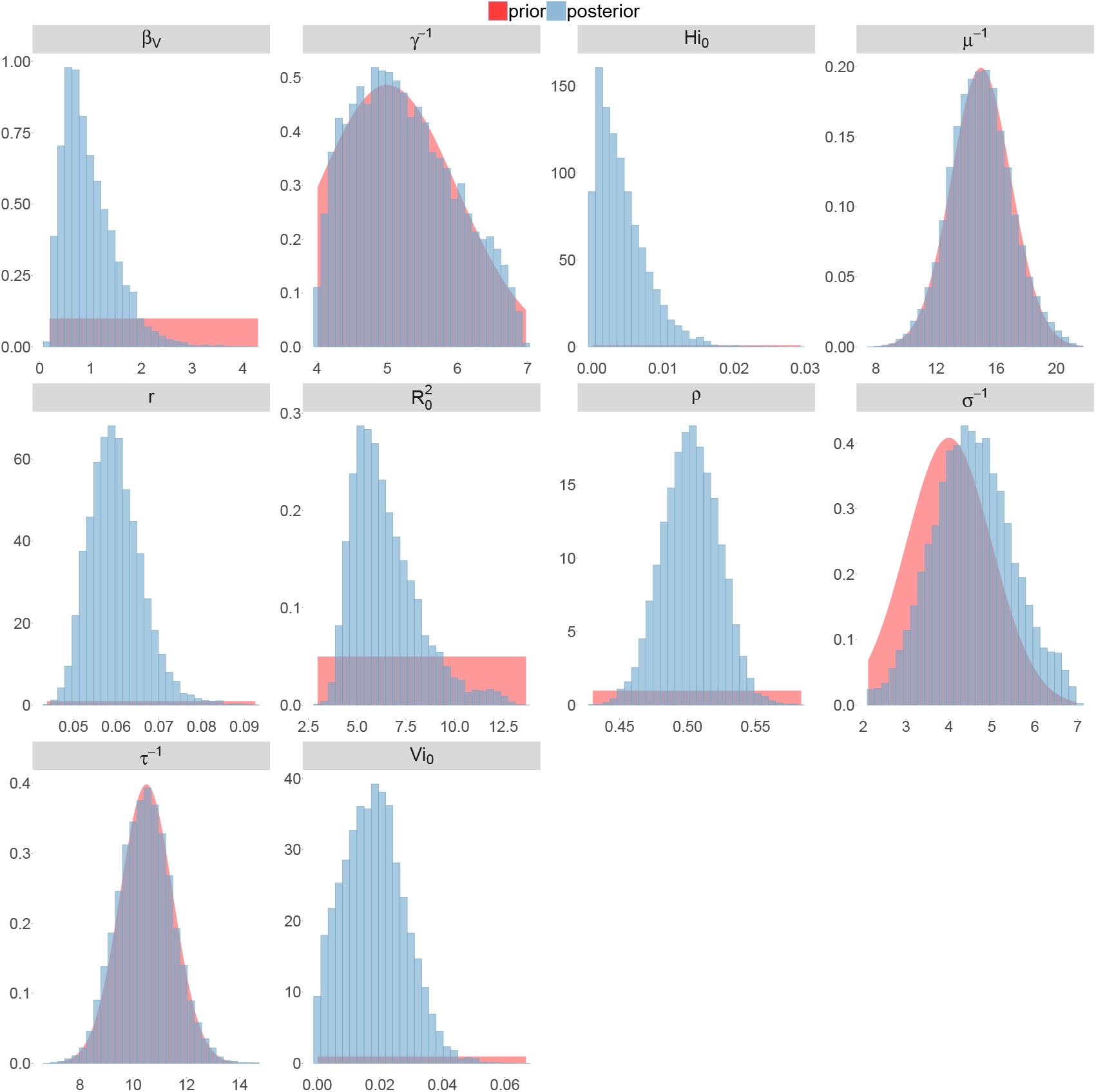
Posterior distributions. Pandey model, Tahiti island.

**Figure 9.**
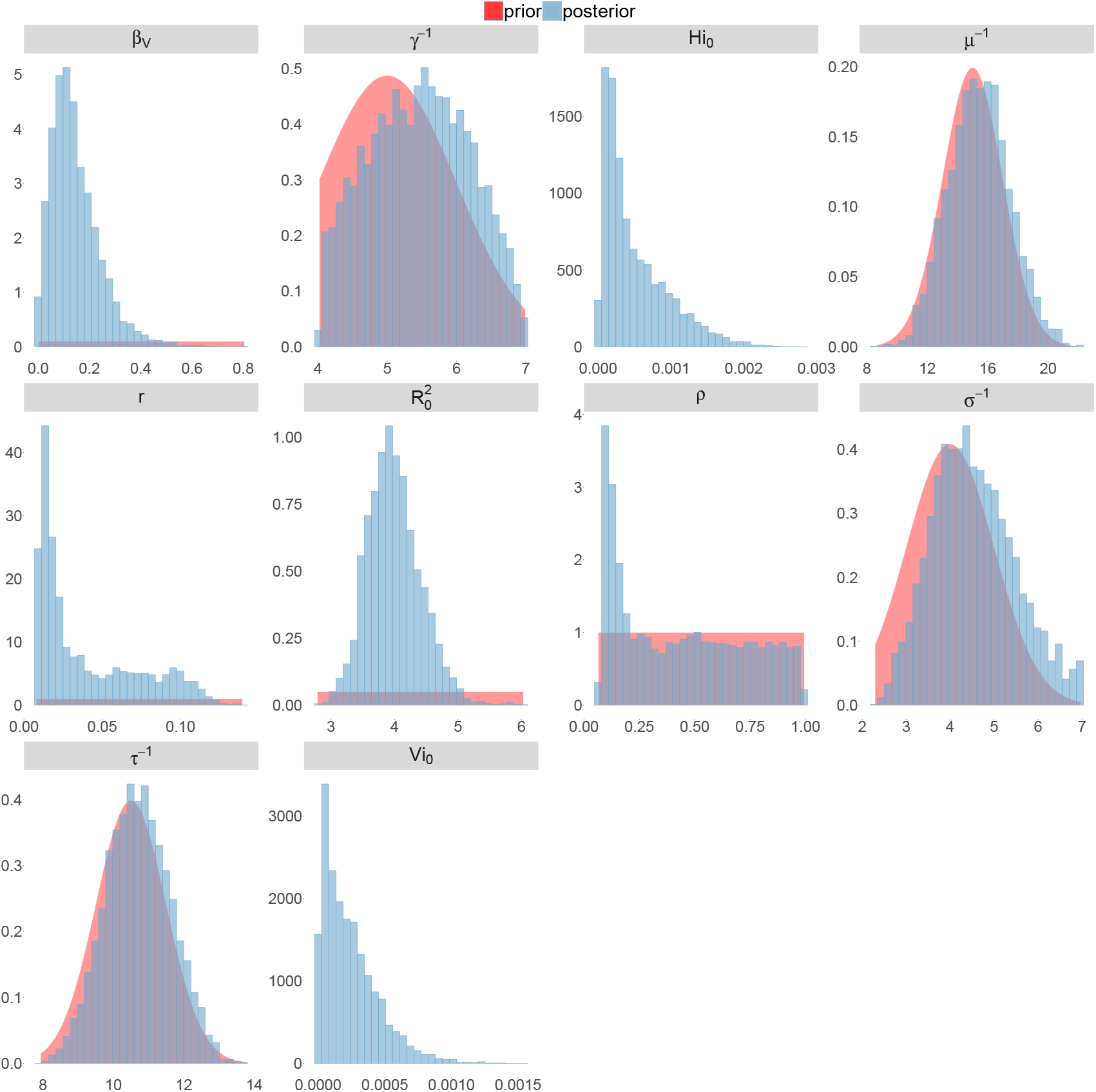
Posterior distributions. Pandey model, New Caledonia.

**Figure 10.**
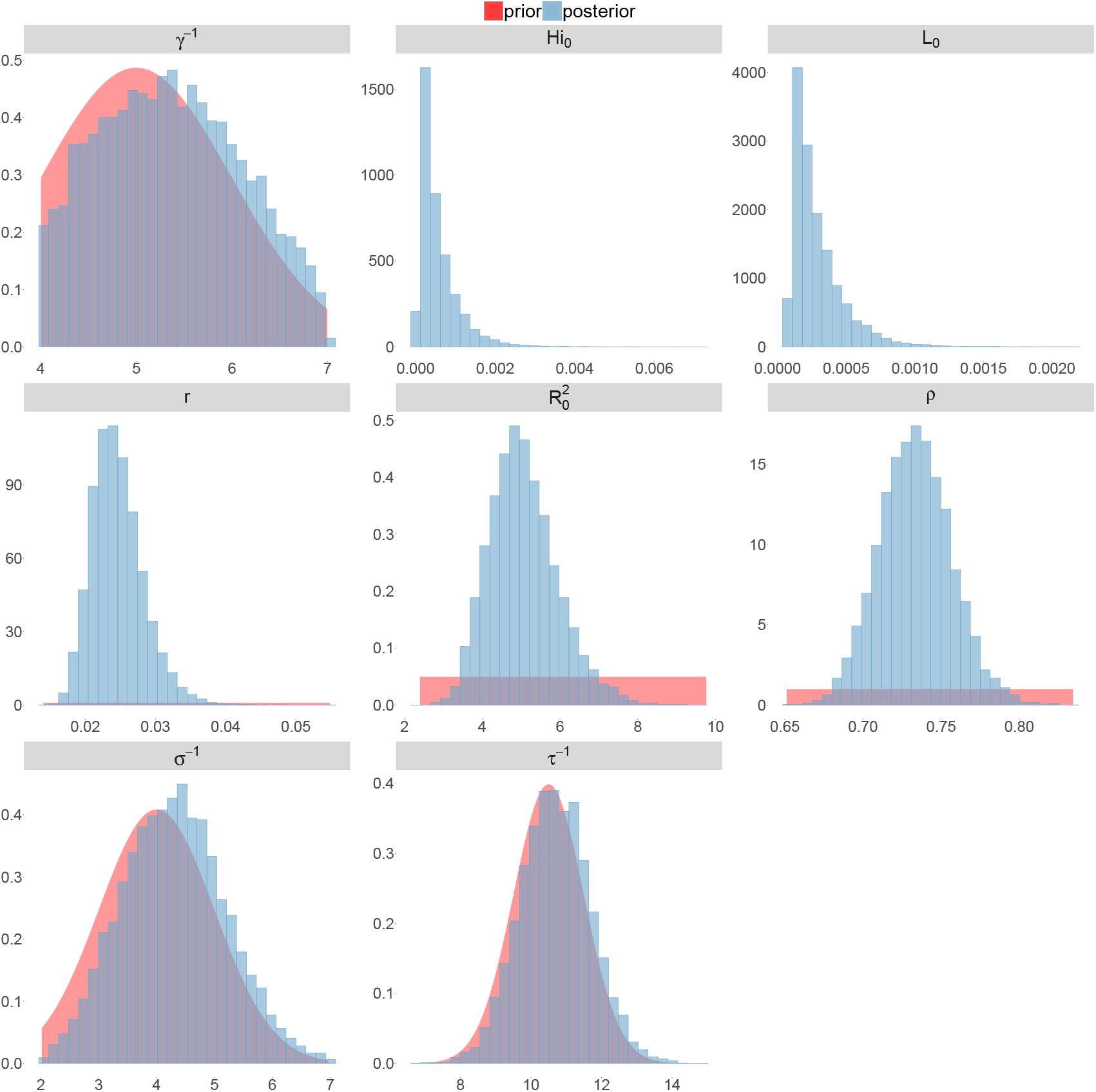
Posterior distributions. Laneri model, Yap island.

**Figure 11.**
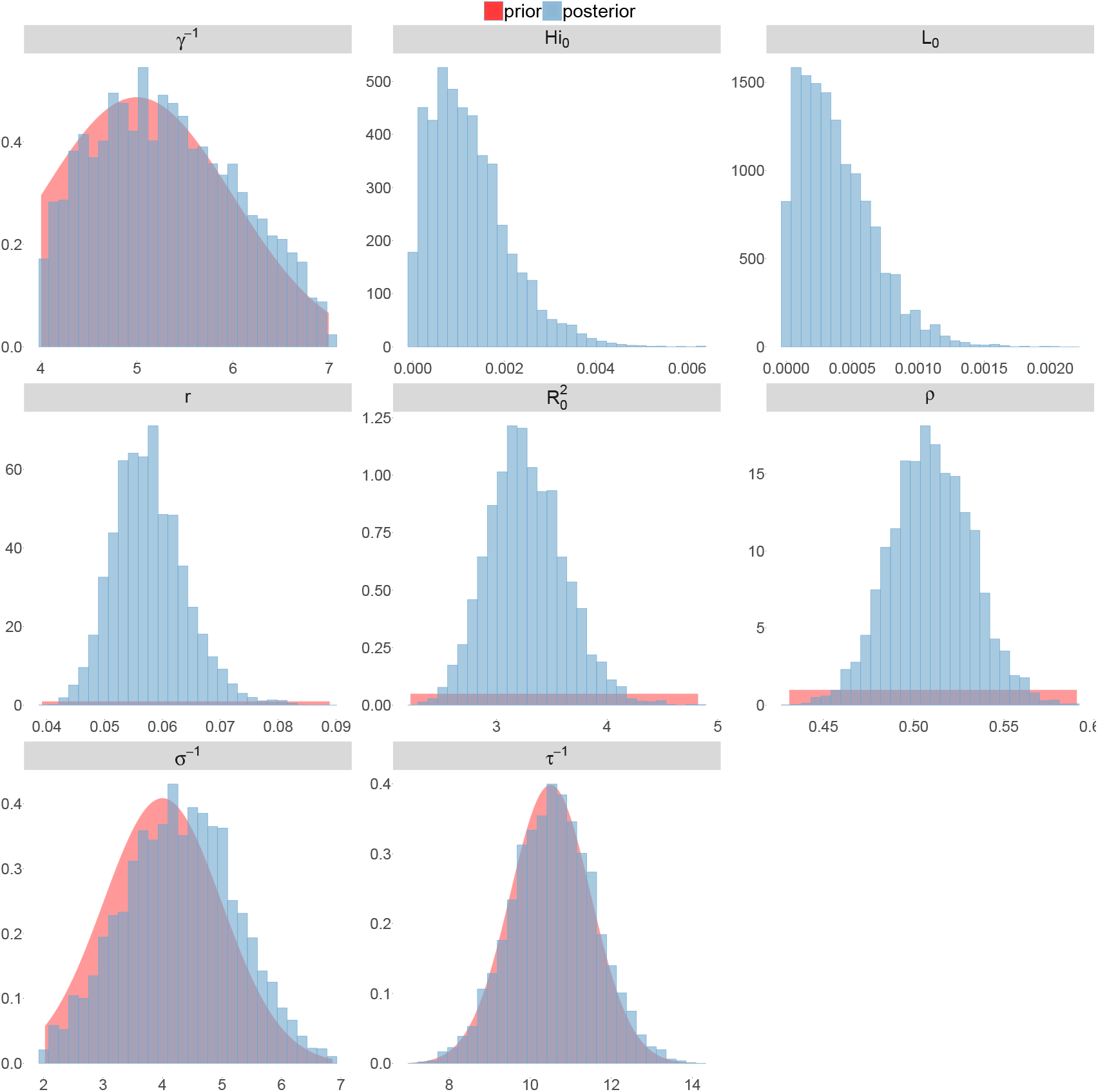
Posterior distributions. Laneri model, Moorea island.

**Figure 12.**
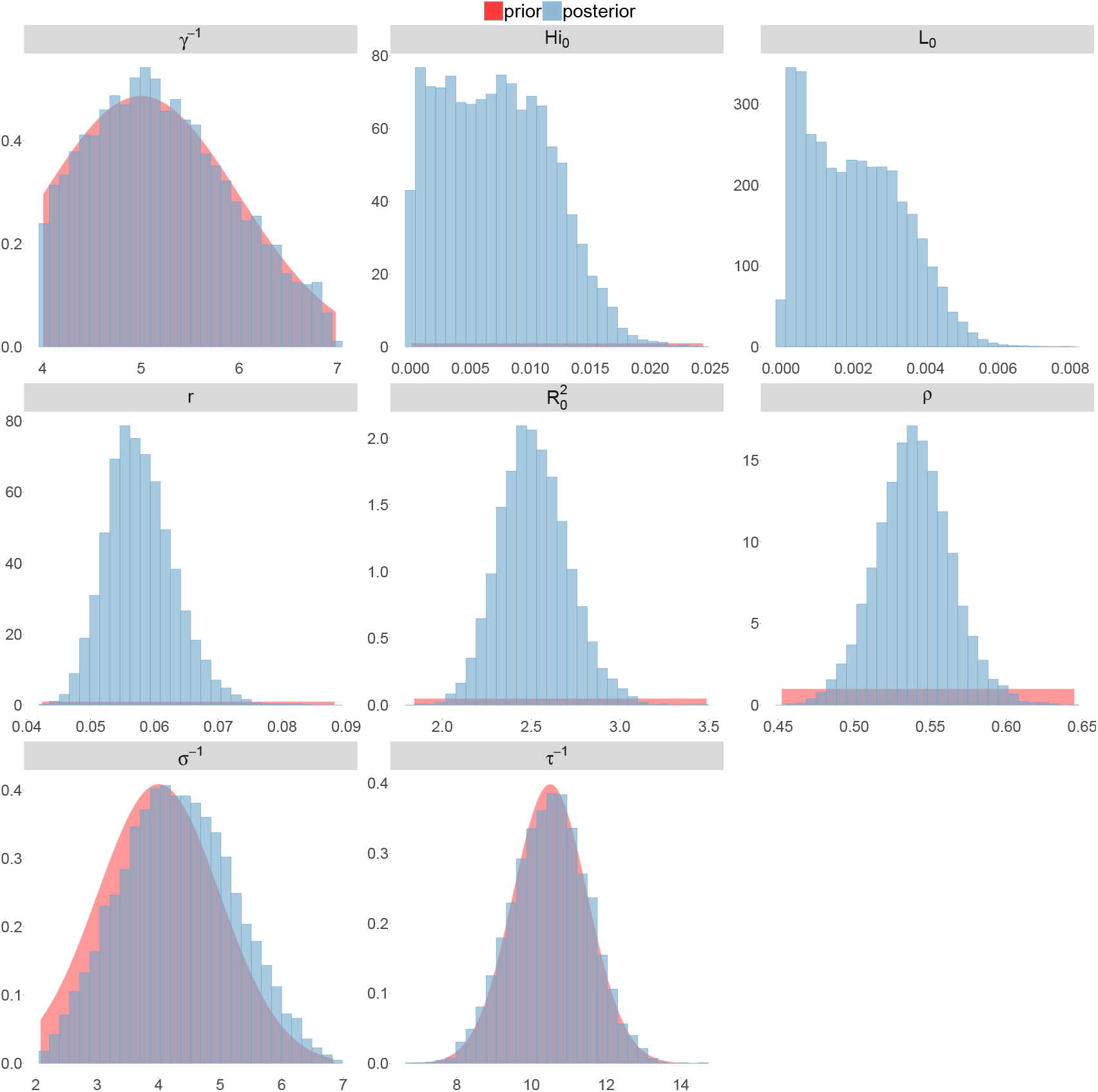
Posterior distributions. Laneri model, Tahiti island.

**Figure 13.**
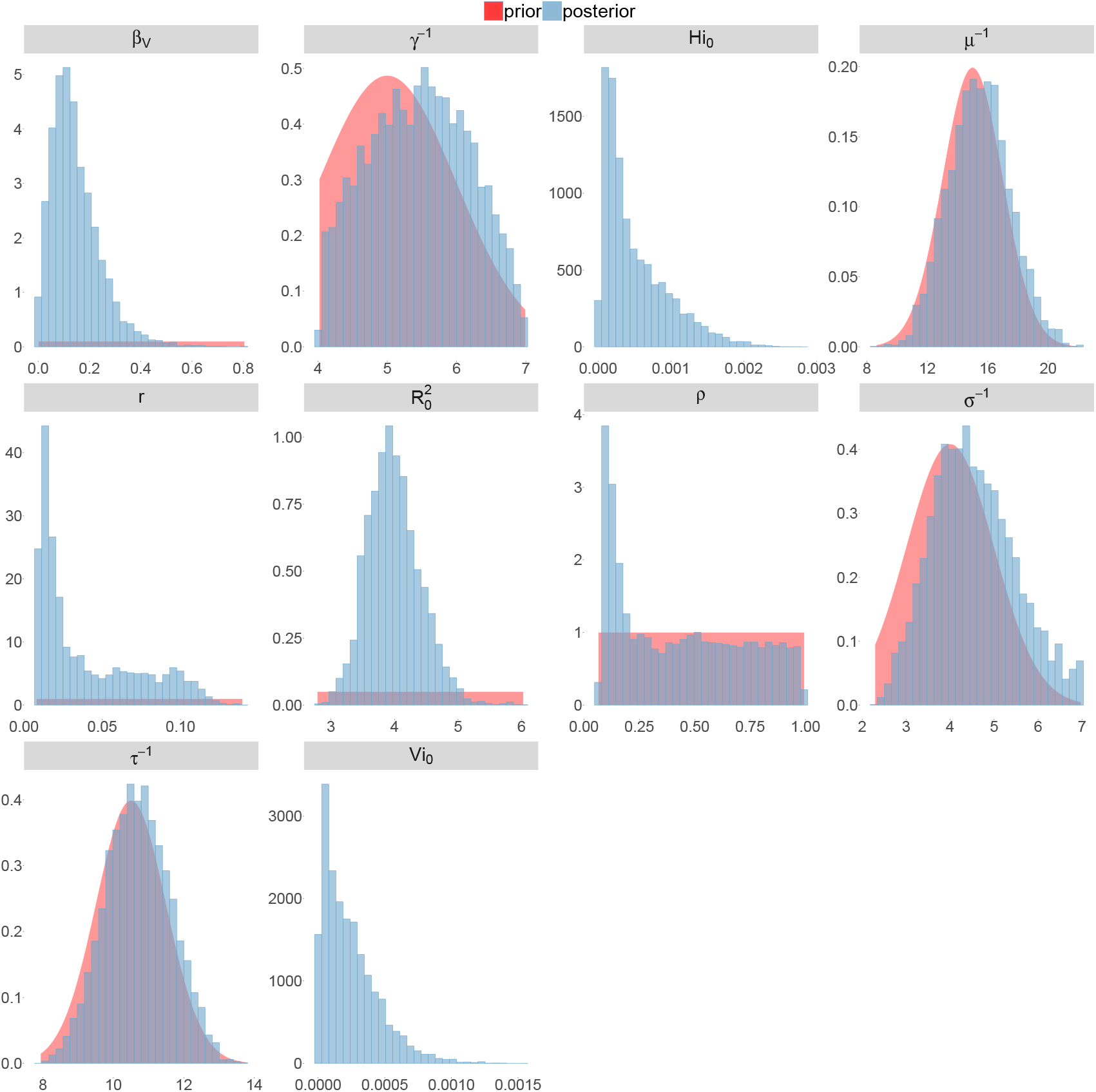
Posterior distributions. Laneri model, New Caledonia.

### Correlation between estimated parameters

The marginal posterior densities (Figures 6-13) do not indicate the correlation between parameters, i.e when the observed value of one parameter is highly influenced by the value of another one. In some cases, this phenomenon reveals identifiability issues: the model manages to estimate only a pair of parameters but cannot identify each one separately. In our case, the observation rate and the fraction of the population involved in the epidemic are strongly negatively correlated when no information is provided on seroprevalence (Figures 9 and 13).

**Figure 14.**
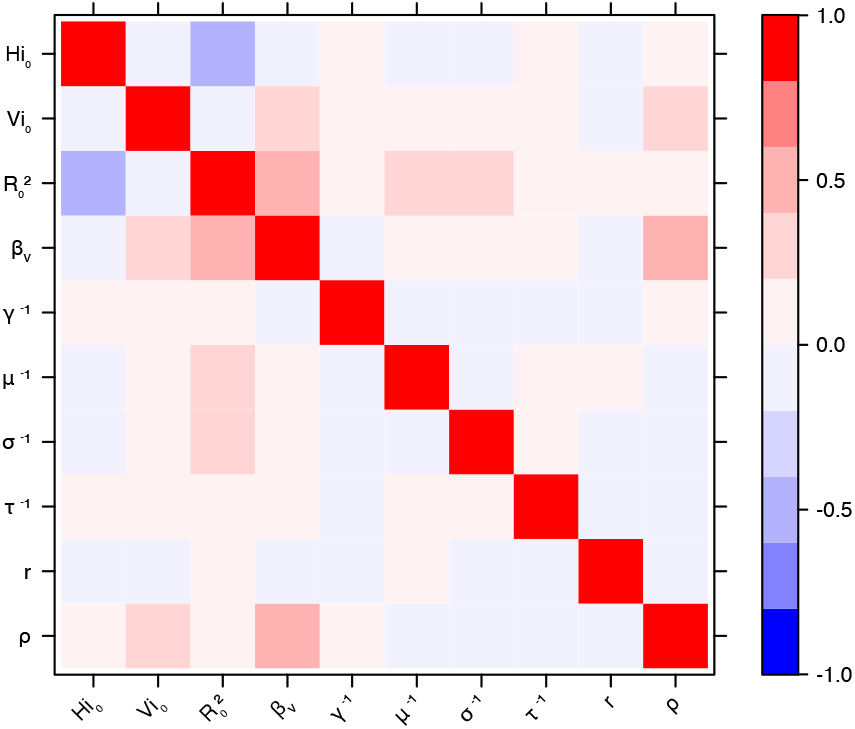
Correlation plot of MCMC output. Pandey model, Yap island.

**Figure 15.**
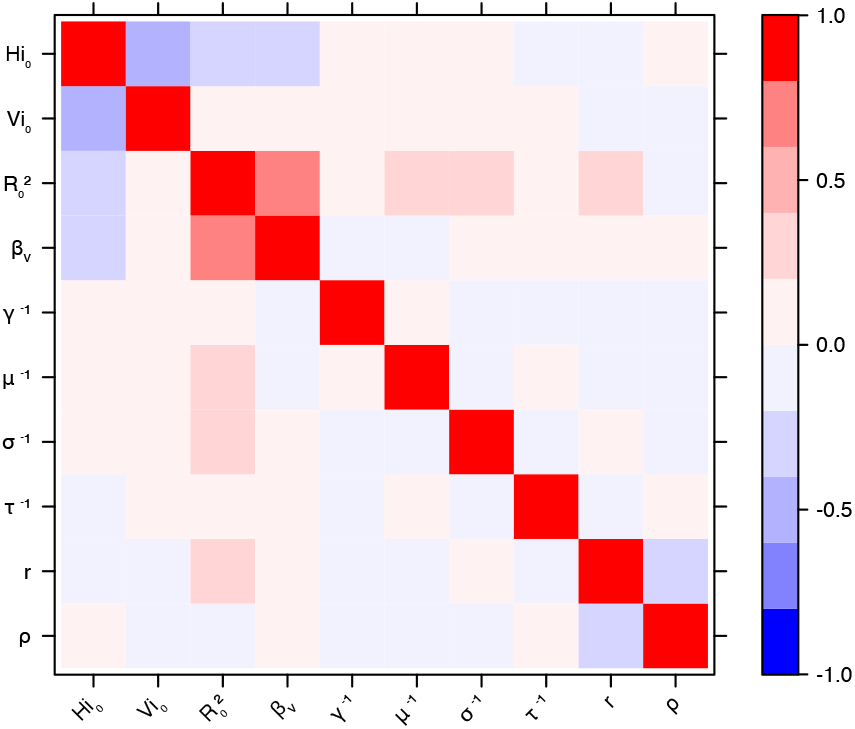
Correlation plot of MCMC output. Pandey model, Moorea island.

**Figure 16.**
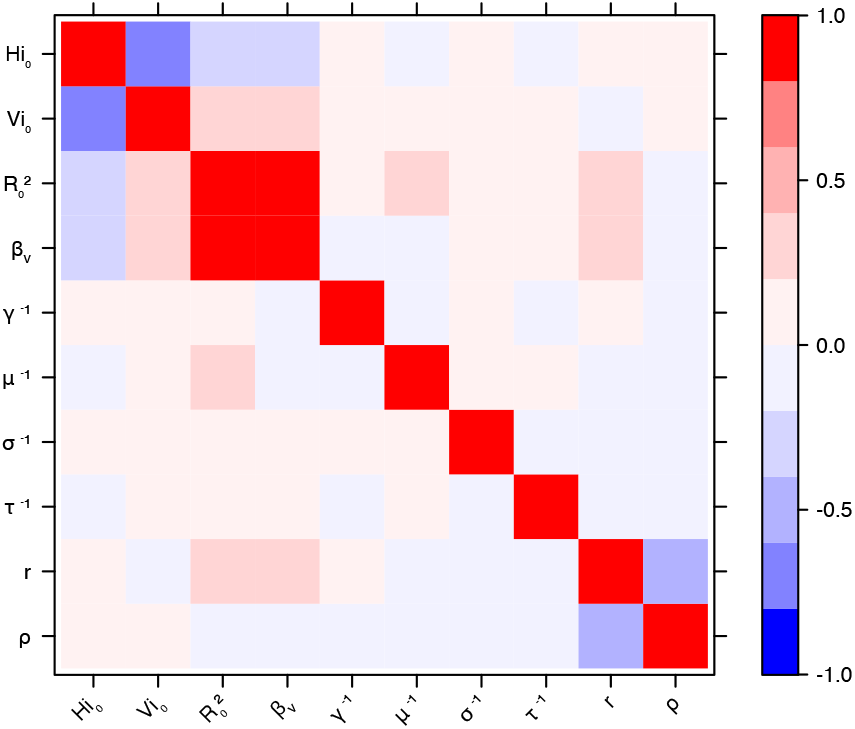
Correlation plot of MCMC output. Pandey model, Tahiti island.

**Figure 17.**
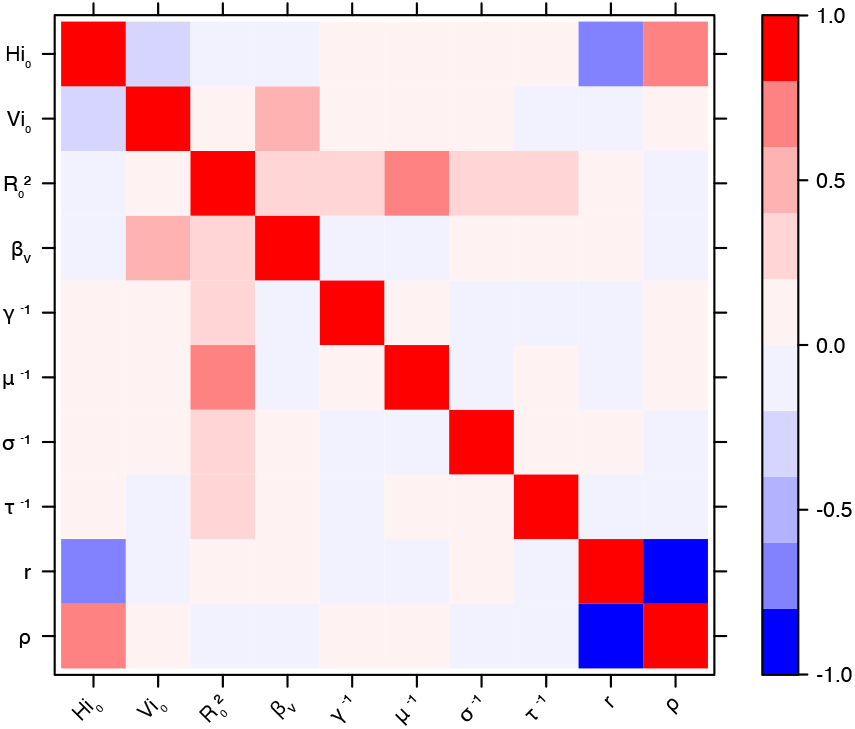
Correlation plot of MCMC output. Pandey model, New Caledonia.

**Figure 18.**
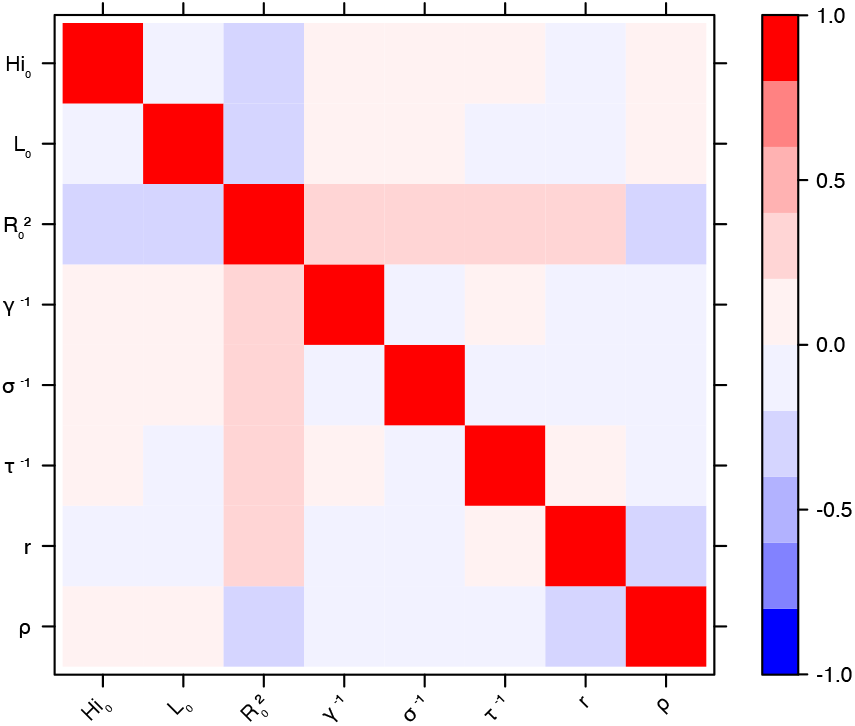
Correlation plot of MCMC output. Laneri model, Yap island.

**Figure 19.**
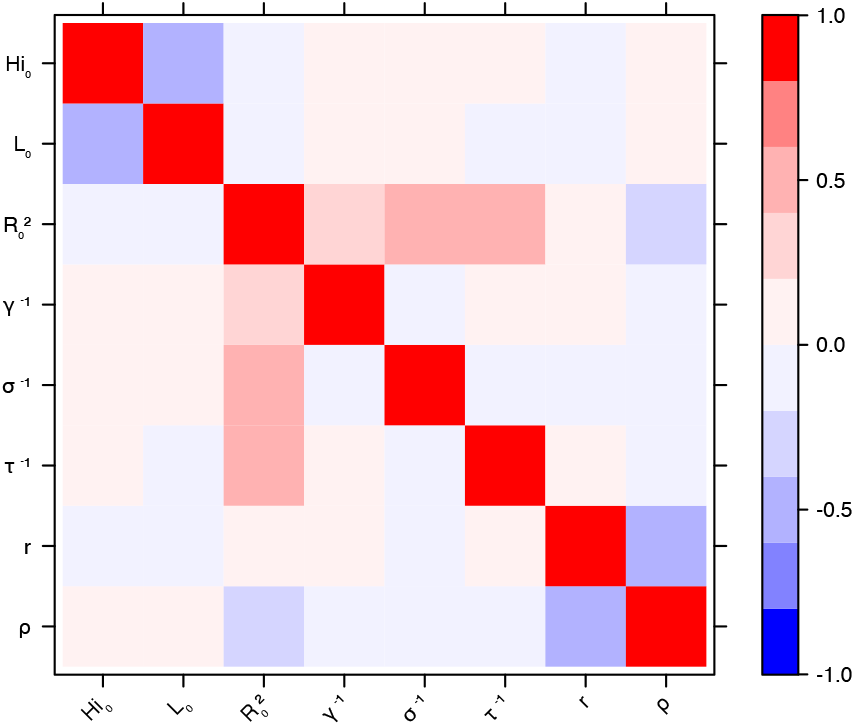
Correlation plot of MCMC output. Laneri model, Moorea island.

**Figure 20.**
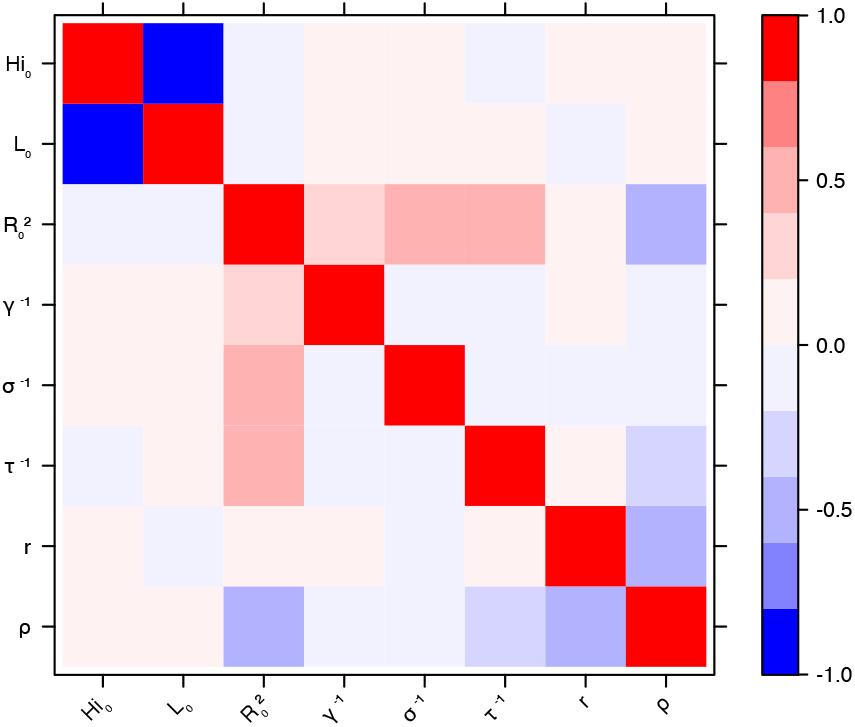
Correlation plot of MCMC output. Laneri model, Tahiti island.

**Figure 21.**
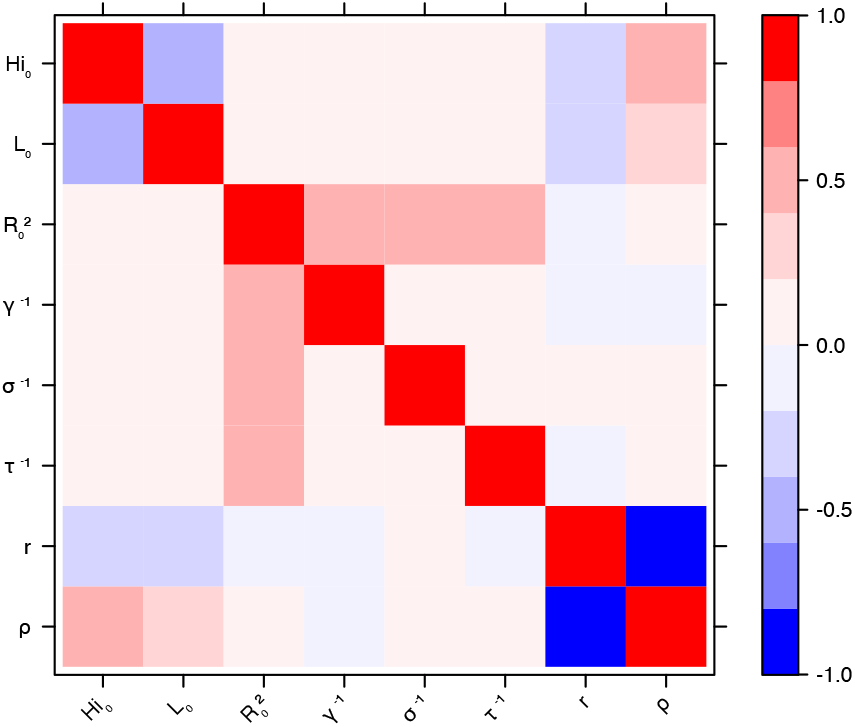
Correlation plot of MCMC output. Laneri model, New Caledonia.

## Acknowledgments

CC, DGS and BC are partially supported by the “Pépiniere interdisciplinaire Eco-Evo-Devo” from the Centre National de la Recherche Scientifique (CNRS). The research leading to these results has also received funding from the European Commission Seventh Framework Program [FP7/ 2007-2013] for the DENFREE project under Grant Agreement n° 282 378. The funders played no role in the study design, data collection, analysis, or preparation of the manuscript.

## Contributors

CC and BC designed the study, CC, DGS and BC contributed to the numerical part of the study, and all the authors participated in the interpretation of the results and in the writing of the manuscript.

